# AAV-mediated ARSA replacement for the treatment of Metachromatic Leukodystrophy

**DOI:** 10.1101/2025.03.12.642609

**Authors:** Shyam Ramachandran, Jeffery Ardinger, Jie Bu, MiAngela Ramos, Lilu Guo, Dhiman Ghosh, Mahmud Hossain, Shih-Ching Chou, Yao Chen, Erik Wischhof, Swathi Ayloo, Roger Trullo, Yuxia Luo, Jessica Hogestyn, Daniel DuBreuil, Emily Crosier, Johanna Adams, Amy Richards, Michael Tsabar, Giorgio Gaglia, Shelley Nass, Bindu Nambiar, Denise Woodcock, Catherine O’Riordan, Qi Tang, Bradford Elmer, Bailin Zhang, Martin Goulet, Christian Mueller

## Abstract

Metachromatic leukodystrophy (MLD) is an autosomal recessive neurodegenerative disorder caused by mutations in the arylsulfatase A (ARSA) gene, resulting in lower sulfatase activity and the toxic accumulation of sulfatides in the central and peripheral nervous system. Children account for 70% of cases and become progressively disabled with death occurring within 10 years of disease onset. Gene therapy approaches to restore ARSA expression via adeno-associated viral vectors (AAV) have been promising but hampered by limited brain biodistribution. We report the development of a novel capsid AAV.GMU01, demonstrating superior biodistribution and transgene expression in the central nervous system of non-human primates (NHPs). Next, we show that AAV.GMU01-ARSA treated MLD mice exhibit persistent, normal levels of sulfatase activity and a concomitant reduction in toxic sulfatides. Treated mice also show a reduction in MLD-associated pathology and auditory dysfunction. Lastly, we demonstrate that treatment with AAV.GMU01-ARSA in NHPs is well-tolerated and results in potentially therapeutic ARSA expression in the brain. In summary, we propose AAV.GMU01-ARSA mediated gene replacement as a clinically viable approach to achieve broad and therapeutic levels of ARSA.

## INTRODUCTION

Metachromatic leukodystrophy (MLD) is a progressive, autosomal recessive neurodegenerative disorder caused by mutations in the *arylsulfatase A (ARSA)* gene, which result in a deficiency of the lysosomal enzyme ARSA and the pathological accumulation of sulfatides in cells (1–3). Over 200 mutations in the *ARSA* gene have been linked to MLD (4), leading to the buildup of 3-O-sulfogalactosyl ceramides (sulfatides) and their deacylated form, lyso-sulfatide, a distinct marker of disease pathology (5, 6). Sulfatide accumulation in oligodendrocytes and Schwann cells (5, 7) drives demyelination in both the central and peripheral nervous systems (CNS and PNS) (8–10), while short-chain fatty acid sulfatides in neurons and astrocytes contribute to neuronal dysfunction, cell loss, and progressive neurodegeneration (8, 11–15). In the CNS, this disruption destabilizes myelin, impairing oligodendrocyte function and leading to widespread demyelination in regions such as the basal ganglia, cerebellum, and spinal cord, resulting in cognitive decline, motor dysfunction, spastic tetraparesis, ataxia, spasms, and seizures (13, 16, 17). Similarly, PNS demyelination causes peripheral neuropathy, muscle weakness, and sensory deficits (13). Sulfatide accumulation also triggers calcium buildup and cellular stress (18), further exacerbating myelin instability and neurodegeneration (19). While non-neural tissues, such as the kidneys and gallbladder, are also affected (20), clinical symptoms are primarily concentrated in the nervous system, making the restoration of ARSA activity in both the CNS and PNS the primary therapeutic goal to mitigate the toxic effects of sulfatide accumulation.

MLD manifests in three primary forms, late-infantile, juvenile, and adult, delineated by the age of onset. While the initial symptoms vary, all patients experience progressive and severe disability, regression of motor skills, gait difficulties, ataxia, and weakness. As the disease progresses, patients experience dysphagia, feeding intolerance, seizures, hypotonia, and peripheral neuropathy. Those with late infantile MLD have a mean life expectancy of around four years (21) Currently, Atidarsagene autotemcel (Libmeldy/Lenmeldy, Orchard Therapeutics), an autologous hematopoetic stem cell therapy, is the sole approved disease-modifying therapy for MLD (17). However, its approval is limited to pre-symptomatic late infantile or early-juvenile patients without neurological impairment (22). Other treatment options are limited to palliative care, symptom alleviation, tube feeding, and psychological support. Thus, a considerable unmet need still exists for MLD patients.

Due to their ability to deliver sustained expression of their transgene cargo, adeno-associated virus (AAV) mediated gene therapy has shown success in treating neurological disorders (23–26). Over 300 patients across 14 types of neurological disorders have undergone AAV infusions showing good tolerance and safety (27–30). Here, we report the development of a novel AAV-mediated gene therapy for MLD. This therapy comprises a novel capsid, AAV-GMU01 with enhanced CNS targeting in nonhuman primates (NHP) and a wild-type human *ARSA* cDNA. We show that delivery of AAV-GMU01-*ARSA* into the CNS of *Arsa* knockout mice results in a reduction of toxic sulfatides and phenotypic reversal. In NHPs, we demonstrate broad biodistribution and a CNS-wide therapeutic footprint—enabled by ARSA protein cross-correction—following delivery via a minimally invasive and clinically feasible administration route. Finally, we establish in NHPs that this treatment delivers potentially therapeutic doses.

## RESULTS

### The novel capsid AAV.GMU01 yields superior biodistribution compared to AAV.rh10 in cynomolgus NHPs

MLD affects all regions of the nervous system, and an effective therapy requires widespread distribution of ARSA protein. AAV.rh10, a naturally occurring capsid, efficiently targets the CNS in both rodents and NHPs (31, 32). However, despite promising pre-clinical results (33–35), a clinical trial employing intra-parenchymal delivery of AAV.rh10 encoding ARSA did not achieve therapeutic endpoints (36). We aimed to identify a capsid which could yield enhanced biodistribution and superior transgene expression when delivered via a minimally invasive, intra-CSF route of administration.

Screening of a 7-amino-acid peptide-insert library based on the AAV9 capsid in Rhesus macaques via intracerebroventricular (ICV) injection identified AAV.GMU01 as a novel capsid with significantly enhanced vector biodistribution and transgene expression in critical CNS regions compared to the parental AAV9 capsid (Application #20240100194). To further evaluate its therapeutic potential, we compared the biodistribution and transgene expression of an eGFP-encoding AAV.GMU01 to AAV.rh10 following intrathecal delivery (**Supplemental Table 1**). We administered AAV.GMU01-CBA-*eGFP* or AAV.rh10-CBA-*eGFP* to cynomolgus monkeys at the cervical level 1-2 junction at a dose of 2.75e13 vector genomes (VG) per animal; 16-29 days later we evaluated vector exposure and eGFP expression. Despite displaying approximately 10-fold lower vector exposure across 41 grey matter punches (representing 16 brain regions) than AAV.rh10-treated (**Figure 1A-C**), AAV-GMU01 exhibited improved eGFP expression (**Figure 1D-F**). To visualize this difference, we plotted the correlation between eGFP expression and vector genomes, either by animal or by each of the 41 grey matter punches. This plot demonstrates a leftward shift in AAV.GMU01 correlation, relative to AAV.rh10, which signifies higher expression at lower brain VG exposure (**Figure 1G-H**). Immunohistochemistry against eGFP confirmed robust expression in AAV.GMU01-treated NHPs (**Figure 1I**). AAV.GMU01-treated NHPs also exhibited lower vector genomes in the spinal cord, dorsal root ganglia (DRGs), liver, heart, lung, and kidney with only the spleen showed higher exposure (**Supplemental Figure 1A**). Despite this, eGFP levels in spinal cord and DRGs were higher than or similar to those achieved by AAV.rh10 (**Supplemental Figure 1B**). These results indicate that AAV.GMU01 achieves widespread CNS biodistribution and transgene expression at a significantly lower AAV vector exposure than AAV.rh10.

**Figure 1.**
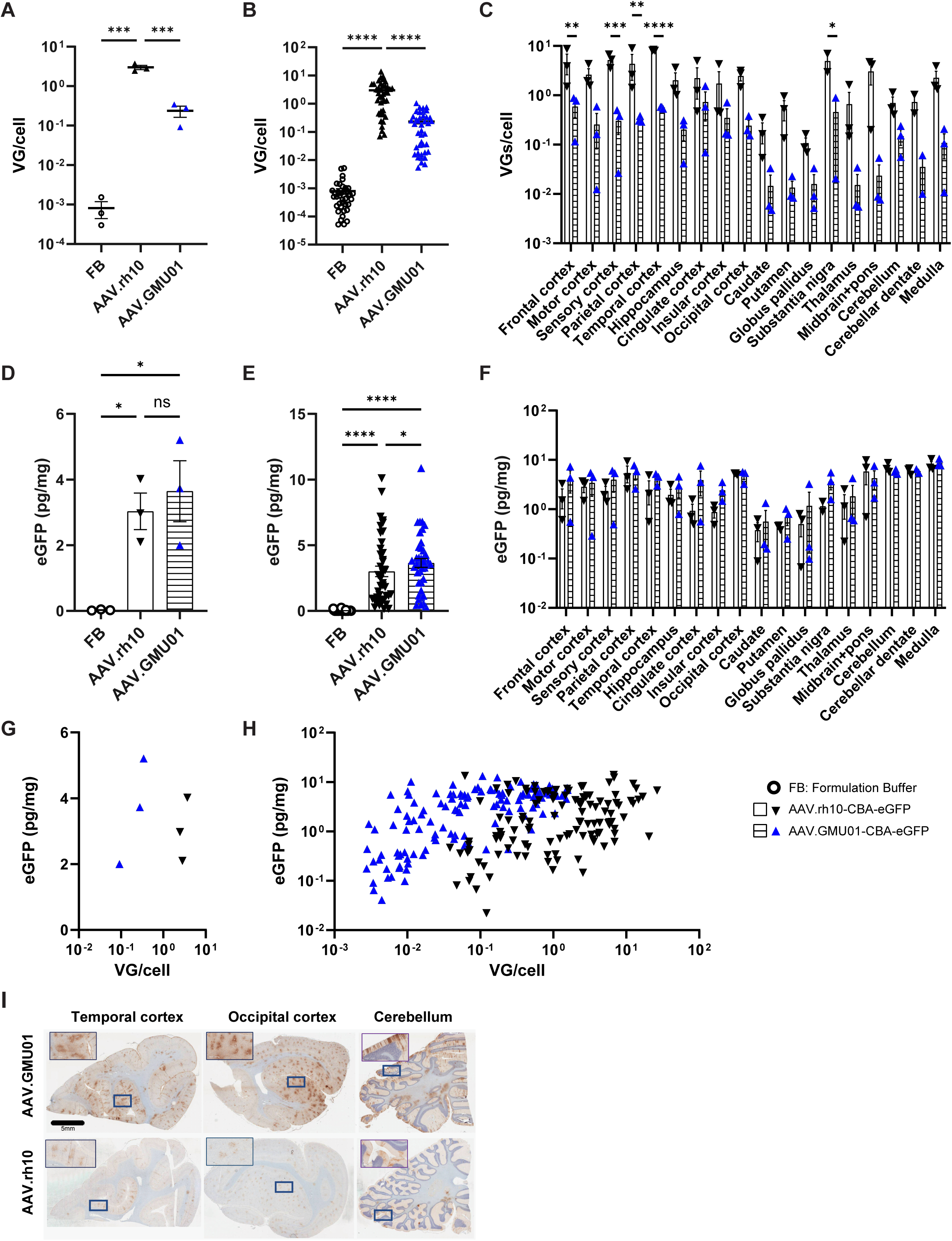
The novel capsid AAV.GMU01 shows higher transgene expression in the brain of NHPs compared to AAV.rh10. Cynomolgus monkeys (Male, Mauritian 2yr old, 2-3kg) seronegative for AAV.rh10 and AAV.GMU01 were dosed intrathecal at the cervical level 1-2 junction using a ported intrathecal catheter inserted at the lumbar region. Animals were dosed in the Trendelenburg position. One dose of AAV.GMU01 or rh10-CBA-eGFP was administrated at 2.75e13VG/NHP (3.65e11VG/gm brain weight). 16-29 days post-dosing, animals were euthanized. (**A-H**) 41 tissue biopsy punches from 19 different gray matter brain regions were taken. AAV vector genome copies (**A-C**) were measured in brain biopsy punches by bGH dPCR and normalized to *TUBB1* gene intron to obtain VG copies per cell, presented by animal (**A**) or by punch (**B, C**). eGFP expression (**D-F**) in the same regions were measured by ELISA, presented by animal (**D**) or by punch (**E, F**). (**G, H**) Correlation of vector exposure to eGFP expression in brain, presented by animal (**G**) or by punch (**H**). (**F**) Representative images depicting distinct brain regions from AAV.GMU01 and AAV.rh10 treated NHPs stained for eGFP. Error bars represent mean with standard error. Two-way ANOVA with Tukey’s or Sidak (panel B, D) multiple comparison test. \**p*<0.05; \*\**p*<0.01; \*\*\**p*<0.001; \*\*\*\**p*<0.0001.

### Comparison and selection of CSF delivery route

While the intravenous (IV) and intraparenchymal routes of administration can achieve AAV distribution to the CNS, both have considerable drawbacks. IV delivery requires high doses which present the risk of toxicity in non-target tissue. Intra-CSF delivery achieves more widespread biodistribution to the CNS at lower doses than IV administration (37–40) and low concentrations of immunoglobulin G in the CSF make intra-CSF delivery less prone to risks related to pre-existing neutralizing antibodies (41). Meanwhile, intraparenchymal delivery often results in non-homogenous distribution (42). Therefore, intra-CSF is the preferred route of administration for CNS-targeted therapies. To establish an optimal intra-CSF route of delivery, we examined the biodistribution of AAV.GMU01 delivered by one of two different routes: 1) Bilateral ICV, and 2) direct injection to the cisterna magna (ICM). We chose to evaluate the ICM route both because it is a clinically feasible and minimally invasive method for direct access to nervous system tissues (37–40, 43), and because it achieves better biodistribution to the CNS than intrathecal-lumbar administration (39, 42, 44, 45).

Cynomolgus NHPs were administered AAV.GMU01-CBA-nLuc-mCherry at a dose of 2.0e13 VG per animal (**Supplemental Table 2**). After four weeks, we collected brain, spinal cord, DRGs, and peripheral tissues, analyzing vector genomes and luciferase activity. Histopathological analysis and quantification of neurofilament light chain (NfL) were also conducted to assess the relative tolerability of each route of administration (RoA). In 20 brain regions, both administration routes demonstrated a broad distribution of vector genomes (**Figure 2A**). Cortical regions exhibited transduction levels of approximately one VG/cell, with ICV delivery showing a trend towards increased transduction in subcortical structures. ICV delivery demonstrated significantly increased NanoLuciferase (nLuc) activity across the brain, particularly in cortical and subcortical regions (**Figure 2B**). Comparable vector biodistribution and transgene activity were observed in the spinal cord, DRGs (**Figure 2C-D**), and other peripheral organs (**Supplemental Figure 2A-B**). Neither RoAs resulted in behavioral signs and both caused a similar increase in NfL in the CSF (**Supplemental Figure 2C**). However, minimal to marked inflammatory and degenerative histopathological findings were noted in the brain of ICV treated NHPs, whereas the ICM group exhibited only minimal findings (**Figure 2E-F**). In the spinal cord and DRGs, minimal to moderate inflammatory and degenerative findings were observed in all groups (**Supplemental Figure 2D-G**), an almost universal sub-clinical finding after AAV gene therapy (46). To evaluate cellular tropism in the NHP brain, we conducted an immunofluorescent co-detection assay, demonstrating transduction to neurons, astrocytes, microglia and oligodendrocytes (**Supplemental Figure 3**). Additional studies using single-nucleus RNA sequencing would be needed to further explore cell-type-specific transduction across NHP brain tissues. In summary, we concluded that ICM delivery of AAV.GMU01 is well-tolerated, and results in widespread transduction in the CNS.

**Figure 2:**
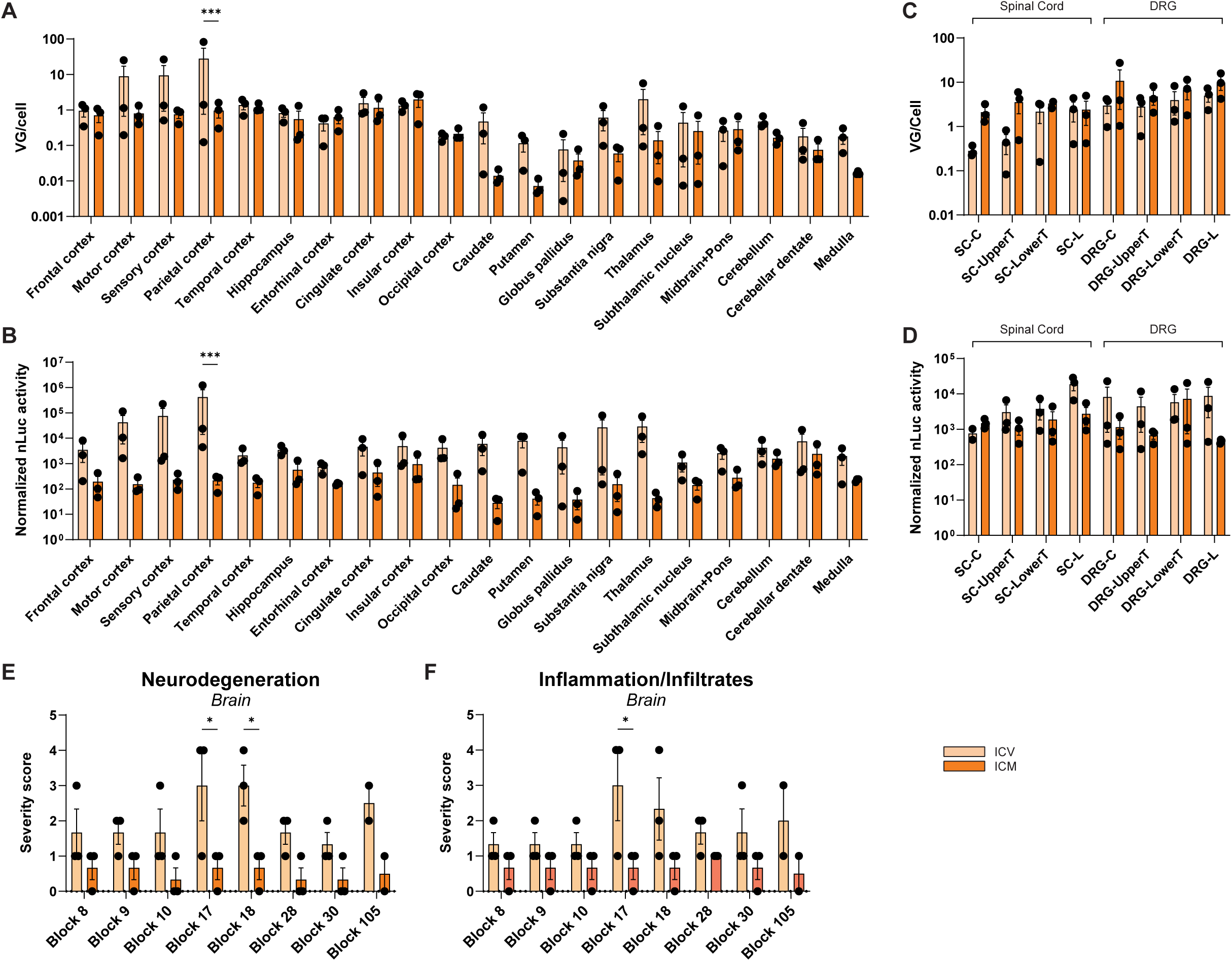
AAV.GMU01 shows widespread vector biodistribution and transgene activity throughout brain, spinal cord and DRGs. Cynomolgus monkeys (Male, Vietnam, 2-3yr old, 2-3kg) seronegative for AAV.GMU01 were dosed with AAV.GMU01-CBA-NLuc-mCherry at 2.0e13VG/NHP (2.75e11 VG/gm brain weight) either by bilateral intracerebroventricular injection (ICV) or direct injection to the cisterna magna (ICM). Four weeks post-dosing, animals were euthanized, and brain, spinal cord, and dorsal root ganglia (DRG) tissues were flash-frozen. Tissue biopsy punches corresponding to the indicated brain regions and 4 spinal cord levels were assessed for vector genome exposure (**A, C**) by dPCR. Nanoluciferease (nLuc) activity (**B, D**) was assessed using the Nano-Glo luciferase assay and normalized to total protein measured by bicinchoninic acid (BCA) assay. (**E**) Histopathological findings in brain, spinal cord, and DRG; each data point represents the maximum severity of findings scored on 1-2 sections per animal. Severity scores refer to findings graded as 0= no findings, 1=minimal, 2=mild, 3=moderate, 4=marked, 5=severe. Block 8: frontal cortex, striatum; Block 9: Insular cortex, temporal cortex; Block 10: motor cortex, striatum; Block 17: thalamus, insular cortex; Block 18: hippocampus, entorhinal cortex; Block 28: parietal cortex; Block 30: cerebellum, medulla, Block 105: midbrain, pons. Error bars represent mean with standard error. One/Two-way ANOVA with Sidak’s multiple comparison test. \**p*<0.05; \*\**p*<0.01; \*\*\**p*<0.001; \*\*\*\**p*<0.0001

### AAV.GMU01-ARSA administration is well tolerated, long lasting and effective in MLD mice

To evaluate the short- and long-term efficacy of ARSA replacement, we conducted four pharmacology studies in MLD mice using AAV.GMU01-*ARSA*, targeting different stages of neuropathological disease progression (**Supplemental Figure 4A**). The MLD mouse model (referred to as *Arsa* KO) was generated by Sanofi (described in supplemental information) and exhibits characteristic sulfatide accumulation in the CNS and visceral organs, evidence of modest gliosis and microglial activation, and an auditory deficit phenotype similar to the original Gieselmann model (47, 48). We selected the ICV administration route for its reproducibility and efficient AAV vector delivery into the CSF of mice. Additionally, in wild-type mice treated with AAV.GMU01-eGFP at a dose of 1.6e11 VG per mouse, single-cell RNA sequencing revealed transduction of approximately 12% of cells in the forebrain and 3% in the hindbrain (**Supplemental Figure 4B-E**).

#### Long-term pharmacology and efficacy

To assess efficacy of long-term ARSA replacement, we administered AAV.GMU01-*ARSA* at a dose of 1.6e11 VG per mouse to pre-neuropathic *Arsa* KO mice and monitored them for 13 months; formulation-buffer treated wild-type (WT) and Arsa KO mice served as controls (Study design: **Supplemental Table 3**). AAV.GMU01 vector genomes and *ARSA* mRNA expression were measured in brain and spinal cord samples (**Supplemental Figure 5A-B**). and sulfatase activity was measured in several brain regions as well as spinal cord, DRGs, sciatic nerve, liver, and plasma. AAV.GMU01-*ARSA* treatment restored sulfatase activity to levels equivalent to WT levels in the brain, spinal cord, DRGs, and sciatic nerve (**Figure 3A**). In the liver and plasma, sulfatase activity was higher in treated mice compared to WT (**Figure 3A-B**). We observed a corresponding reduction in sulfatide deposition, which is the primary driver of toxicity in patients. Treatment normalized lyso-sulfatide (lyso-ST), C16, C18 and total ST levels in the brain and spinal cord and significantly reduced lyso-ST, C16, C18 and total ST levels in DRGs and the sciatic nerve (**Figure 3C**, **Supplemental Figure 6A-C**). MLD patients exhibit an increase in sulfatides in biological fluids (5). making this a critical biomarker readout in the clinic. We observed a substantial reduction in total sulfatides in plasma and cerebrospinal fluid of treated mice (**Figure 3D**). In the brain and spinal cord, the decrease in sulfatide levels coincided with a significant reduction in neuroinflammatory marker levels (*Gfap* and *Aif1*) and in the expression of *Lamp1*, a marker of immune cell activation and degradative lysosomes (**Figure 3E-G**). Furthermore, compared to buffer treated *Arsa* KO mice, AAV.GMU01-*ARSA* treated mice displayed a significant reduction in plasma neurofilament light chain (Nf-L) levels. (**Supplemental Figure 6D**).

**Figure 3:**
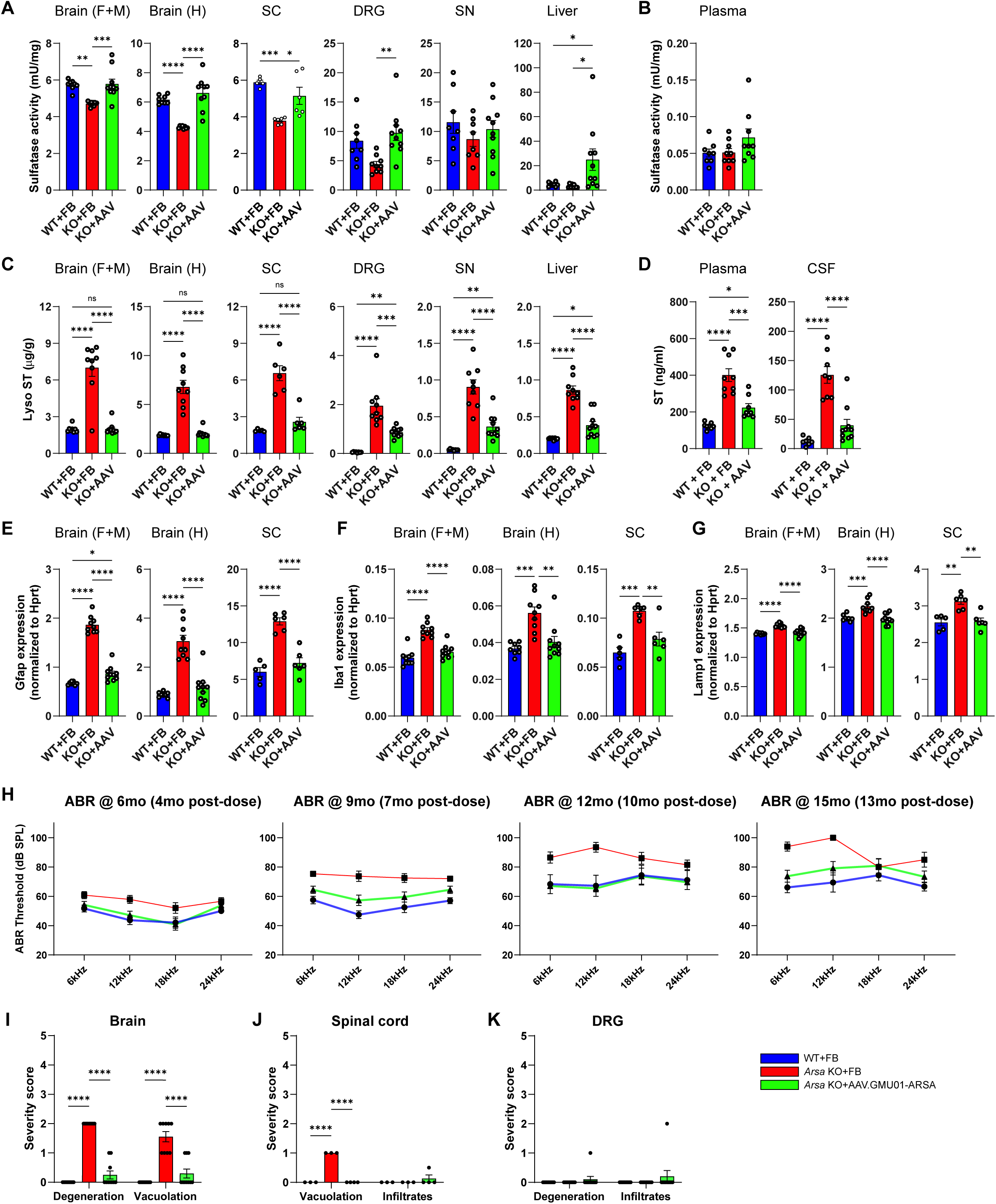
Phenotypic reversal in *Arsa* KO mice treated with AAV.GMU01-*ARSA*. Pre-neuronopathic *Arsa* KO mice and age-matched control animals were dosed with AAV.GMU01-*ARSA* at 1.6e11VG/mouse (3.3e11VG/gram brain weight). Thirteen months post-dose, *Arsa* KO mice were euthanized and brain, spinal cord, DRG, sciatic nerve, liver, plasma, and CSF samples collected. (**A-B**) ARSA-mediated sulfatase activity was measured using the Sulfatase Activity Assay; data normalized to total protein measured by bicinchoninic acid (BCA) assay. (**C-D**) Sulfatide levels were measured using liquid chromatography–mass spectrometry (LC-MS). Data normalized to tissue weight: converted from ng/mL (50 μL) to μg/g (ng ST/μg total protein for DRG and SN). For fluids, data is presented as ng/mL (CSF). **(E-G)** RT-dPCR was performed to quantify *Gfap, Aif1* (gene for Iba1) and *Lamp1* levels, normalized to mouse *Hprt* gene. Each data point represents a single animal. Error bars represent mean with standard error; one-way ANOVA with Tukey’s multiple comparison test. (**H**) At 4-, 7-, 10- and 13-months post-dose, ABR measurements were recorded via electrodes placed on the scalp of an anesthetized animal. Error bars represent mean with standard error; Two-way ANOVA with Tukey’s multiple comparison test. # denote “KO+FB” vs “KO+AAV” group; * denote “WT+FB” vs “KO+FB” group. (**I-K**) Histopathological findings in brain, spinal cord, and DRG; each data point represents the maximum severity of findings scored on 1-2 sections per animal. Severity scores refer to findings graded as 0= no findings, 1=minimal, 2=mild, 3=moderate, 4=marked, 5=severe. Error bars represent mean with standard error. \**p*<0.05; \*\**p*<0.01; \*\*\**p*<0.001; \*\*\*\**p*<0.0001. FB= Formulation Buffer, sulfatide= ST.

Hearing impairment is common in MLD patients. The Auditory Brainstem Response (ABR) test provides functional insights into both the inner ear (cochlea) and central hearing pathways. Mice and humans perceive multiharmonic communication sounds similarly (49) with mice having best hearing range between 12 and 20kHz. While mice perceive high-frequency sounds (12-20kHz) better than low-frequency (3-12kHz), like humans, they respond better to low-frequency sounds (49). *Arsa* KO mice exhibit a progressive decline in sound perception over time, with the largest change in ABR observed at lower frequencies (**Figure 3H**). Treatment with AAV.GMU01-*ARSA* largely prevented the loss of hearing in *Arsa* KO mice (**Figure 3H**) with the difference between groups appearing as early as 4 months post-dosing (earliest time point assessed), indicating the potentially rapid physiological and functional benefits of AAV-mediated ARSA gene replacement. Consistent with other models, *Arsa* KO mice displayed histopathological changes, including vacuolation and neurodegeneration, chiefly in the brainstem and/or cerebellum (**Figure 3I**). Treated mice exhibited a notable reduction in both the severity and incidence of these histological changes (**Figure 3I**). Additionally, all untreated *Arsa* KO mice exhibited vacuolation in the spinal cord which was absent in both wild-type and treated mice (**Figure 3J**). Histopathological alterations in the dorsal root ganglia occurred in only one treated mouse (**Figure 3K**).

#### Longitudinal pharmacology

This study aimed to assess target engagement and reversibility of MLD-associated pathological and biochemical traits in the *Arsa* KO model over a 6-month period. Six-month-old early-neuronopathic *Arsa* KO mice underwent bilateral ICV injections of AAV.GMU01-ARSA at a dose of 5.0e10 VG per mouse. Control groups, including age-matched WT and *Arsa* KO mice, received doses of formulation buffer (Study design: **Supplemental Table 4**). Animals were monitored for 6 months, with assessments of vector genomes, sulfatase activity, and sulfatide levels at 1-, 2-, 3-, and 6-months post-dose. AAV.GMU01-*ARSA* vector genome exposure in the brain (forebrain + midbrain) remained consistent over the 6-month period (**Figure 4A**). Sulfatase activity was restored to levels equivalent to WT animals (**Figure 4B**) and reached its peak by 1-month post-dose. We observed significant reductions in sulfatide deposition including in lyso-ST, C16, C18 and total ST at all timepoints. (**Figure 4C-F**). Clearance of sulfatides was progressive, returning to normal by 6 months post-dose. Due to low vector exposure in the hindbrain and spinal cord (**Supplemental Figure 7A, G**), we measured only a minimal increase in sulfatase activity; (**Supplemental Figure 7B, H**) however, lyso-ST, C16, C18 and total ST were all reduced in both tissues (**Supplemental Figure 7 C-E, I-K**), suggesting ARSA protein distribution to the hindbrain and spinal cord. Furthermore, AAV.GMU01-*ARSA* treatment significantly reduced total ST levels in cerebrospinal fluid and plasma (**Figure 4G-H**). AAV administration resulted in a marginal increase in plasma neurofilament light chain (NfL) levels, which resolved over time and by six months post dose, plasma NfL levels were significantly decreased (**Figure 4I**).

**Figure 4:**
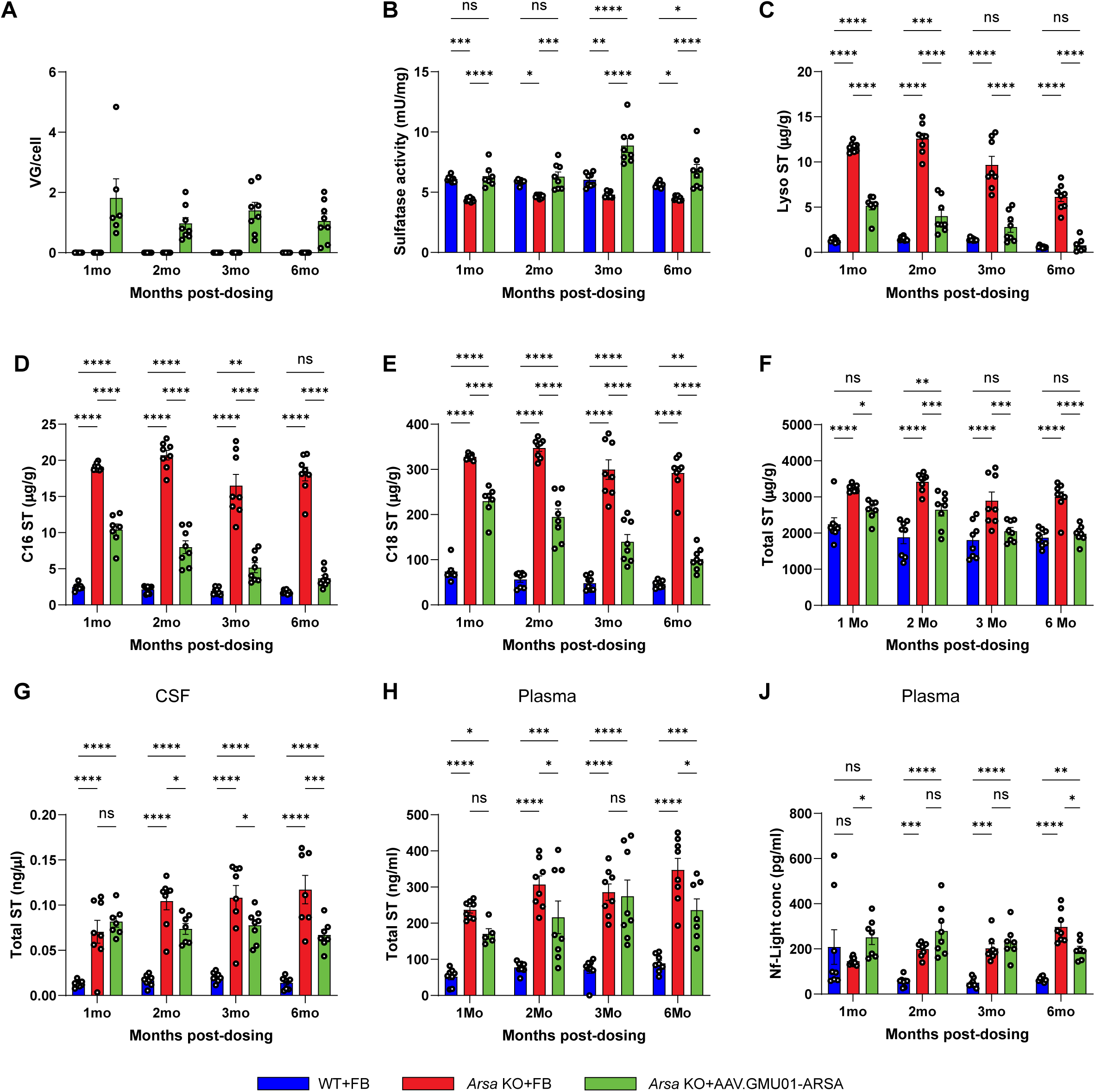
ARSA expression and function is persistent over time. Early-neuronopathic *Arsa* KO mice (6mo at dosing) and age-matched control animals were dosed with AAV.GMU01-*ARSA* at 5e10VG/mouse (1.0e11VG/gram brain weight). One, two, three, and six months post-dose, *Arsa* KO mice were euthanized, and samples collected. (**A-F**) Brain (forebrain + midbrain) samples. (**A**) Vector exposure by bGH-dPCR normalized to the *Rab1a* gene (intronic region). (**B**) ARSA-mediated sulfatase activity was measured using the Sulfatase Activity Assay Kit, data normalized to total protein measured by BCA assay. (**C**) Lyso-sulfatide, (**D**) C16-sulfatide isoform, (**E**) C18-sulfatide isoform and (**F**) Total sulfatide levels were measured using LC-MS. Data normalized to tissue weight: converted from ng/mL (50 μL) to μg/g. Total sulfatide levels were also measured in (**G**) CSF and (**H**) plasma samples. **(I)** Plasma samples were collected at necropsy and were assayed by Quanterix Simoa to quantify neurofilament light chain (NfL). Error bars represent mean with standard error. Two-way ANOVA with Tukey’s multiple comparison test. \**p*<0.05; \*\**p*<0.01; \*\*\**p*<0.001; \*\*\*\**p*<0.0001. FB= Formulation Buffer, sulfatide= ST.

Notably, a reduction in the incidence and severity of neuronal and neuropil changes was evident in the brains of AAV.GMU01-*ARSA* treated *Arsa* KO mice, as early as two months post-dose (**Supplemental Figure 8**). In the brains of some AAV.GMU01-*ARSA* treated mice, perivascular infiltrates of mononuclear cells with minimal severity were observed, usually near the hippocampus (**Supplemental Figure 8**). Additionally, increased cellularity of glial cells and/or mononuclear cells, typically of minimal or occasionally mild severity was also present in the DRGs of certain AAV.GMU01-*ARSA* treated mice. (**Supplemental Figure 8**). No test-article related histopathological were detected in the spinal cord (**Supplemental Figure 8**). Histological changes in the sciatic nerve were observed in most *Arsa* KO mice, with the incidence and severity occasionally increased to minimal or mild in AAV.GMU01-*ARSA* treated *Arsa* KO mice (**Supplemental Figure 8**).

#### Dose-dependent pharmacology

To evaluate dose-dependent ARSA expression and its impact on efficacy (sulfatide clearance) we ran two studies, one in neonatal (dosing at P0) and another in early-neuronopathic (dosing at 6 months of age) *Arsa* KO mice. Mice underwent bilateral ICV injections of AAV.GMU01-*ARSA* at four increasing doses ranging from 1e10VG to 3.3e11VG/gm brain weight. Control groups, including age-matched WT and *Arsa* KO mice, received formulation buffer (Study design: **Supplemental Table 5**). Six months post-dose, animals treated neonatally showed dose-dependent vector biodistribution (**Figure 5A**) and a robust increase in sulfatase activity in the brain (forebrain + midbrain), with levels surpassing those of WT mice at all doses (**Figure 5B**). We also observed a concomitant decrease in sulfatide deposits in the brain (**Figure 5C-D, Supplemental Figure 9A-B**). Similarly, three months post-dose, early-neuronopathic mice showed a robust increase in brain (forebrain + midbrain) sulfatase activity at the top two doses (**Figure 5F**), with a significant decrease in sulfatide deposits in the brain across all doses (**Figure 5G-H, Supplemental Figure 9C-D**).

**Figure 5.**
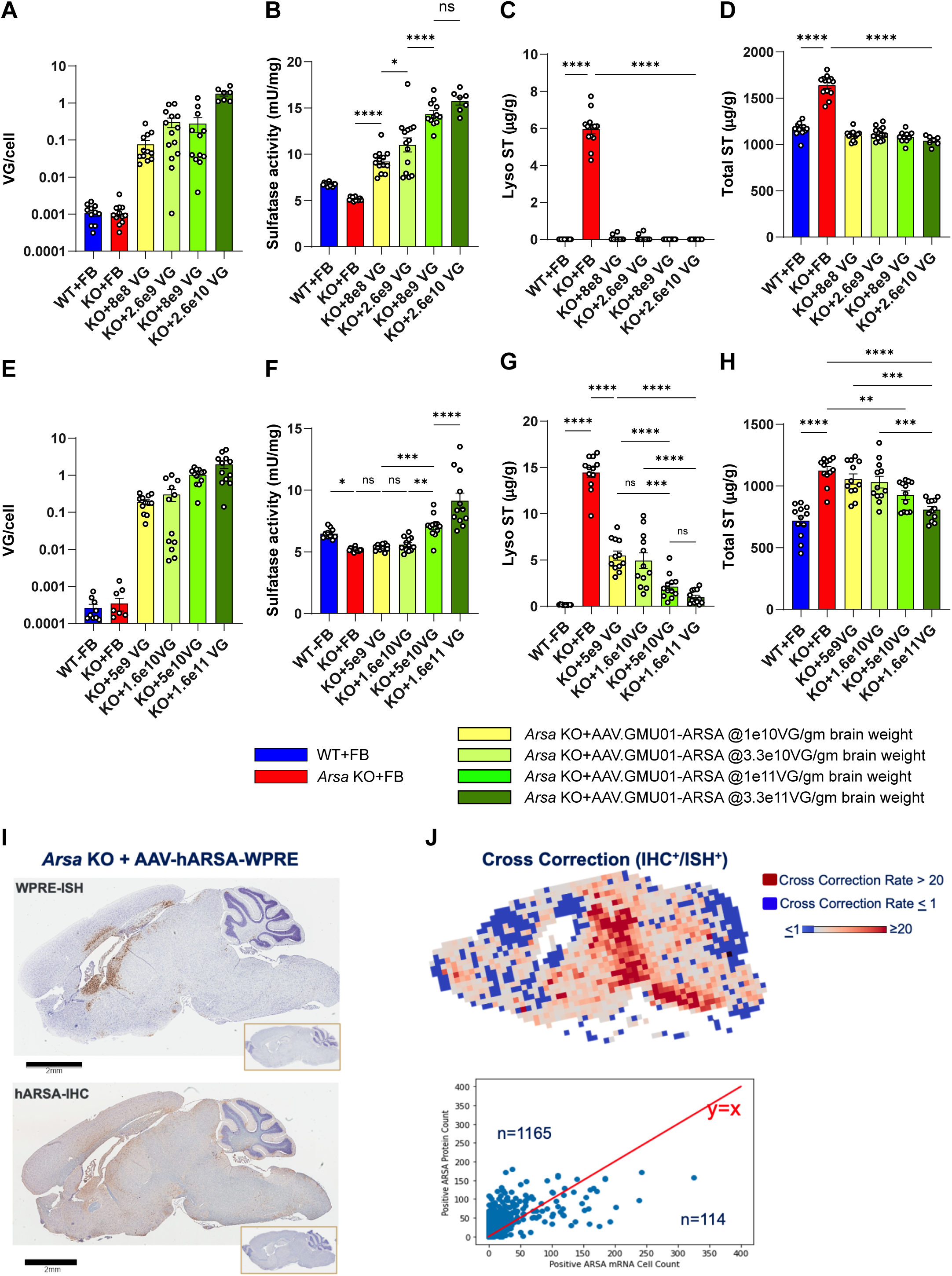
AAV-*ARSA* treatment results in dose-dependent sulfatide clearance in *Arsa* KO mice with evidence of robust cross correction. (A-H) Brain (forebrain + midbrain) samples. (A-D) Neonatal *Arsa* KO mice (P0 at dosing) and age-matched control animals were dosed with AAV.GMU01-*ARSA* at noted doses. Six months post-dose, mice were euthanized and samples collected. (A) Vector exposure by bGH-dPCR normalized to the *Rab1a* gene (intronic region). (B) ARSA-mediated sulfatase activity was measured using the Sulfatase Activity Assay Kit, data normalized to total protein measured by BCA assay. (C) Lyso-sulfatide and (D) Total sulfatide levels were measured using LC-MS. Data normalized to tissue weight: converted from ng/mL (50 μL) to μg/g. (E-H) Early-neuronopathic *Arsa* KO mice (6mo at dosing) and age-matched control animals were dosed with AAV.GMU01-*ARSA* at noted doses. Three months post-dose, mice were euthanized and samples collected. (E) Vector exposure by bGH-dPCR normalized to the *Rab1a* gene (intronic region). (F) ARSA-mediated sulfatase activity was measured using the Sulfatase Activity Assay Kit, data normalized to total protein measured by BCA assay. (G) Lyso-sulfatide and (H) Total sulfatide levels were measured using LC-MS. Data normalized to tissue weight: converted from ng/mL (50 μL) to μg/g. Error bars represent mean with standard error. Two-way ANOVA with Tukey’s multiple comparison test. \**p*<0.05; \*\**p*<0.01; \*\*\**p*<0.001; \*\*\*\**p*<0.0001. FB= Formulation Buffer, sulfatide= ST; gbw= grams per brain weight. (I-J) AAV.rh10-ARSA-WPRE treated *Arsa* KO mice show evidence of ARSA protein cross-correction. Late-stage (13 month) *Arsa* KO mice were dosed with AAV.rh10-CBA-ARSA-WPRE. Three months post-dosing, ARSA-mRNA *in situ* hybridization (ISH) and ARSA-protein immunohistochemistry (IHC) were performed on matched sagittal brain hemi-sections. The sections were imaged and analyzed for signal overlay. (I) Representative sections from mouse brain stained for *ARSA* mRNA (ISH) and ARSA Protein (IHC) with DAPI staining for nuclei (inset: staining from buffer-treated *Arsa* KO mice). (J) Cross-correction factor (ratio of IHC+ cells to ISH+ cells) is represented as a heat map with highly cross-corrected tiles shown in shades of red. ISH+ cell count vs IHC+ cell count from each tile is plotted as a scatter plot with y=x line shown in red. Tiles above the y=x line indicate cross-corrected cells.

Taken together, these data demonstrate that, in *Arsa* KO mice, AAV.GMU01-*ARSA* mediated gene replacement is well tolerated and rectifies MLD-associated phenotypes, validating our therapeutic approach.

### ARSA protein cross-correction generates a large therapeutic footprint

As mentioned above, in certain tissues we observed a discrepancy between sulfatase activity and sulfatide levels with a minimal increase in activity generating a larger than expected reduction in sulfatides. While the relative sensitivity of the two assays likely explains much of this discrepancy, in some tissues (hindbrain, spinal cord) very low vector exposure leads to robust sulfatide clearance. ARSA enzyme is naturally secreted into the extracellular matrix where it is taken up by neighboring and distant cells via the mannose 6-phosphate receptor (M6PR) (50–52). Our data suggests that the recombinant human protein can leverage normal ARSA secretion and reuptake mechanisms facilitating enzyme spread to secondary, non-transduced cells resulting in “cross-correction” in a broad range of cells (50–52) including oligodendrocytes (13, 53) and microglia (54), which are not efficiently transduced *in vivo* by AAV vectors.

To measure the extent of cross-correction, thirteen-month-old *Arsa* KO mice underwent bilateral ICV injections of AAV.rh10-*ARSA* at a dose of 1.6e11 VG per mouse. Three months post-dose, consecutive 5mm brain sections were processed for *ARSA*-mRNA *in situ* hybridization (ISH) and ARSA-protein immunohistochemistry (IHC) (**Figure 5I**), imaged, registered, and the ratio of cells containing ARSA protein to cells containing the vector genome was determined (**Figure 5J**). We observed robust ARSA protein biodistribution throughout the brain, with >84% of brain by area showing evidence of cross-correction (**Figure 5J, Supplemental Figure 10**). In neonatal *Arsa* KO mice dosed bilaterally (ICV) at postnatal day 0 (P0) with AAV.GMU01-*ARSA* at 3.3e11VG/gm brain weight, we performed an immunofluorescent co-detection assay to evaluate cross-correction. These animals demonstrate a robust increase in sulfatase activity and a concomitant decrease in sulfatide deposits in the brain (**Figure 5B-D, Supplemental Figure 9A-B**). Additionally, cross-correction was evident in neurons, astrocytes, and microglia (**Supplemental Figure 11**), highlighting the widespread therapeutic potential of the vector.

### AAV.GMU01-ARSA treatment is well tolerated in NHPs and results in widespread ARSA expression in CNS

To validate our therapeutic approach in a large animal model, we examined the biodistribution and tolerability of AAV.GMU01-*ARSA* in a NHP model. AAV.GMU01-*ARSA* was delivered via ICM infusion to 2-3 year old cynomolgus NHPs at four doses ranging from 7.5e11 to 2.5e13 VG/animal (Study design: **Supplemental Table 6**). Neurological and behavioral tests were performed pre-dose, 7-days post dose, and at necropsy (day 35). Tissue, CSF, and plasma were collected for analysis of VG, *ARSA* mRNA and ARSA protein. A dose dependent increase in AAV.GMU01-*ARSA* vector biodistribution was observed throughout the brain (**Figure 6A**) and grey and white matter punches collected from 19 and 7 brain regions, respectively (**Figure 6B**). This correlated with dose dependent increases in *ARSA* mRNA (**Figure 6C-D**) and protein levels (**Figure 6E-F**) at the 7.5e12 and 2.5e13 VG/animal doses. Additionally, uniform, dose-dependent vector biodistribution (**Supplemental Figure 12A**) and ARSA expression (**Supplemental Figure 12B**) was observed in DRGs and spinal cord along the rostral-caudal axis. We also noted a dose-dependent increase in vector biodistribution in liver, spleen, and cervical lymph node (**Supplemental Figure 12C**).

**Figure 6:**
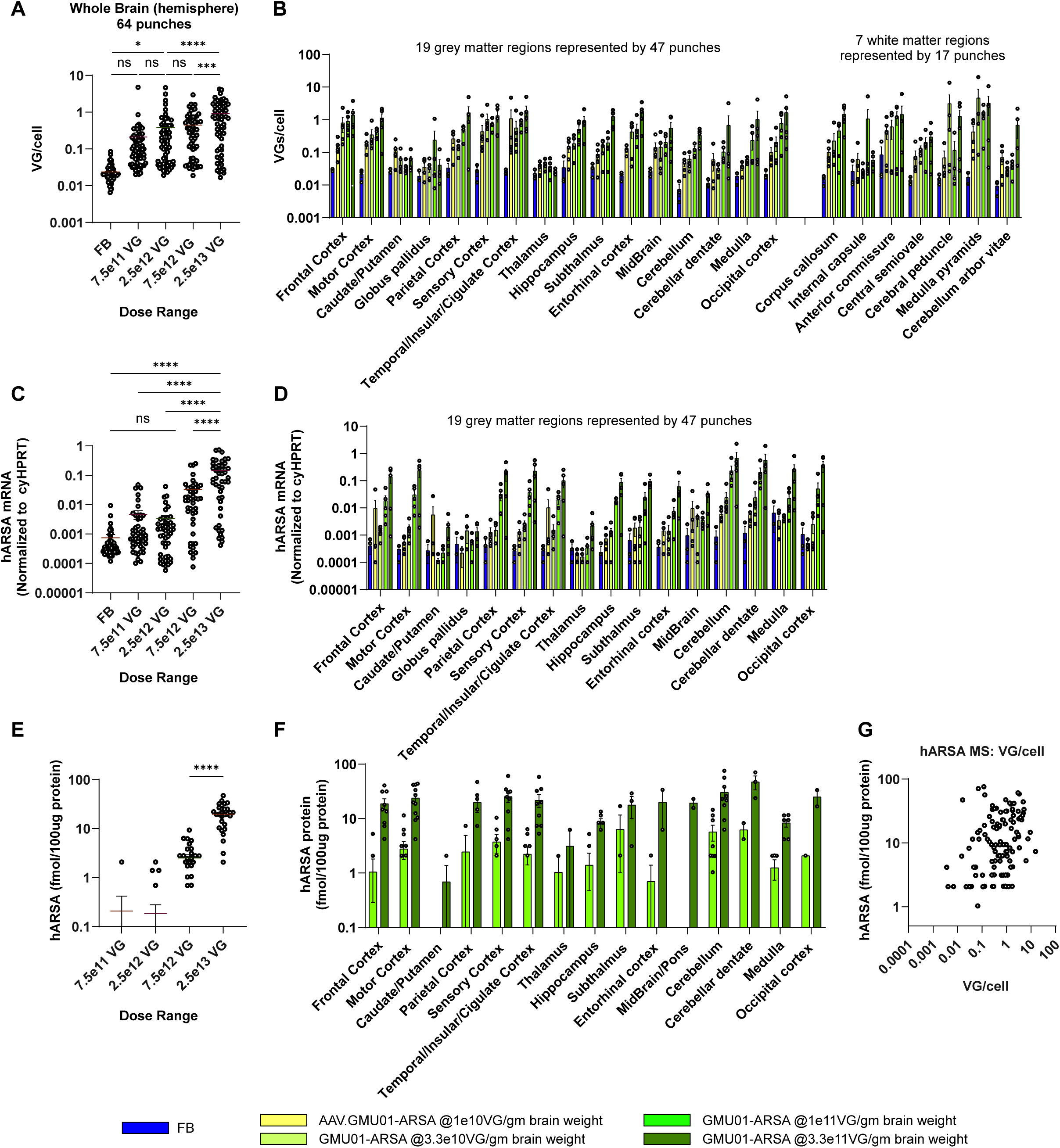
Widespread dose-dependent vector biodistribution and ARSA expression in NHP brain. Purpose bred, naive, male/female cynomolgus (Cambodia 2 to 3 years old, 2.6 to 3.1 kg) NHPs seronegative for AAV.GMU01 neutralizing antibodies were dosed by single direct ICM infusion. Animals were received a single 2.5 mL infusion of AAV.GMU01-*ARSA* at 0.125 mL/min, followed by a 250 uL flush with formulation buffer. Five weeks post-dosing, the animals were euthanized, and samples were assessed. (**A-B**) Vector biodistribution (dPCR) normalized to the *TUBB1* gene intron. Each data point represents VG/cell exposure for that tissue punch, either (**A**) averaged across all NHPs in that group or (**B**) presented by individual brain regions. (**C-D**) *ARSA* mRNA (RT-dPCR) normalized to endogenous *HPRT* gene. Each data point represents normalized ARSA expression in that tissue punch, either (**C**) averaged across all NHPs in that group or (**D**) presented by individual brain regions. (**E-F**) Human ARSA protein expression (LC-MS). Each data point represents the amount of human ARSA in that tissue punch, average across three animals in that group, either presented as an (**E**) average across all NHPs in that group, or (**F**) presented per individual brain regions. Data below LLOQ have been excluded. (**G**) Correlation of ARSA protein and VG/cell at 7.5e12 and 2.5e13VG/NHP doses (R^2^: 0.06, 0.18 respectively). Samples represent 64 biopsy punches from brain representing 19 distinct grey matter regions and 7 distinct white matter regions. Error bars represent mean with standard error. Two-way ANOVA with Tukey’s multiple comparison test. \**p*<0.05; \*\**p*<0.01; \*\*\**p*<0.001; \*\*\*\**p*<0.0001. FB= Formulation Buffer; gbw= grams per brain weight.

Clinical signs (functional or behavioral deficits) were absent in NHPs at 1 or 5 weeks (necropsy) post-dosing in either dosing groups (**Supplemental Table 7**). ICM infusion led to an increase in CSF Nf-L levels, which was more pronounced in the AAV.GMU01-*ARSA* treated animals; the increase was not dose dependent (**Figure 7A**). No significant change in plasma cytokine concentrations were observed at any dose (**Figure 7B, Supplemental Figure 13**), and IFN-γ ELISpot indicated the absence of cell-mediated immune responses to either the GMU01 capsid or ARSA protein at all doses (**Figure 7C**). Sub-clinical microscopic findings were observed in the brain, DRG, spinal cord and peripheral nerves (**Figure 7D-F, Supplemental Figure 14**), including scant degenerate neurons surrounded by gliosis present in the cortex, cerebellar Purkinje cells and rarely in the thalamus (**Figure 7D, Supplemental Figure 14A**). Neuronal degeneration in the spinal cord consisted of nerve fiber degeneration, while in peripheral nerves, degeneration phenotype of the sciatic, femoral, and radial nerves was also observed (**Figure 7E, Supplemental Figure 14B-C**). Neuronal degeneration in the DRG was noted at all test-article doses (**Figure 7F, Supplemental Figure 14D-E**). No test article related findings were observed in visceral organs (heart, liver, gallbladder, spleen, pancreas, adrenal gland, lung, bone, sternum/marrow, ovary, duodenum, testis, epididymis, thymus, eye, uterus with cervix, kidney) at any dose (**Supplemental Figure 14F**).

**Figure 7:**
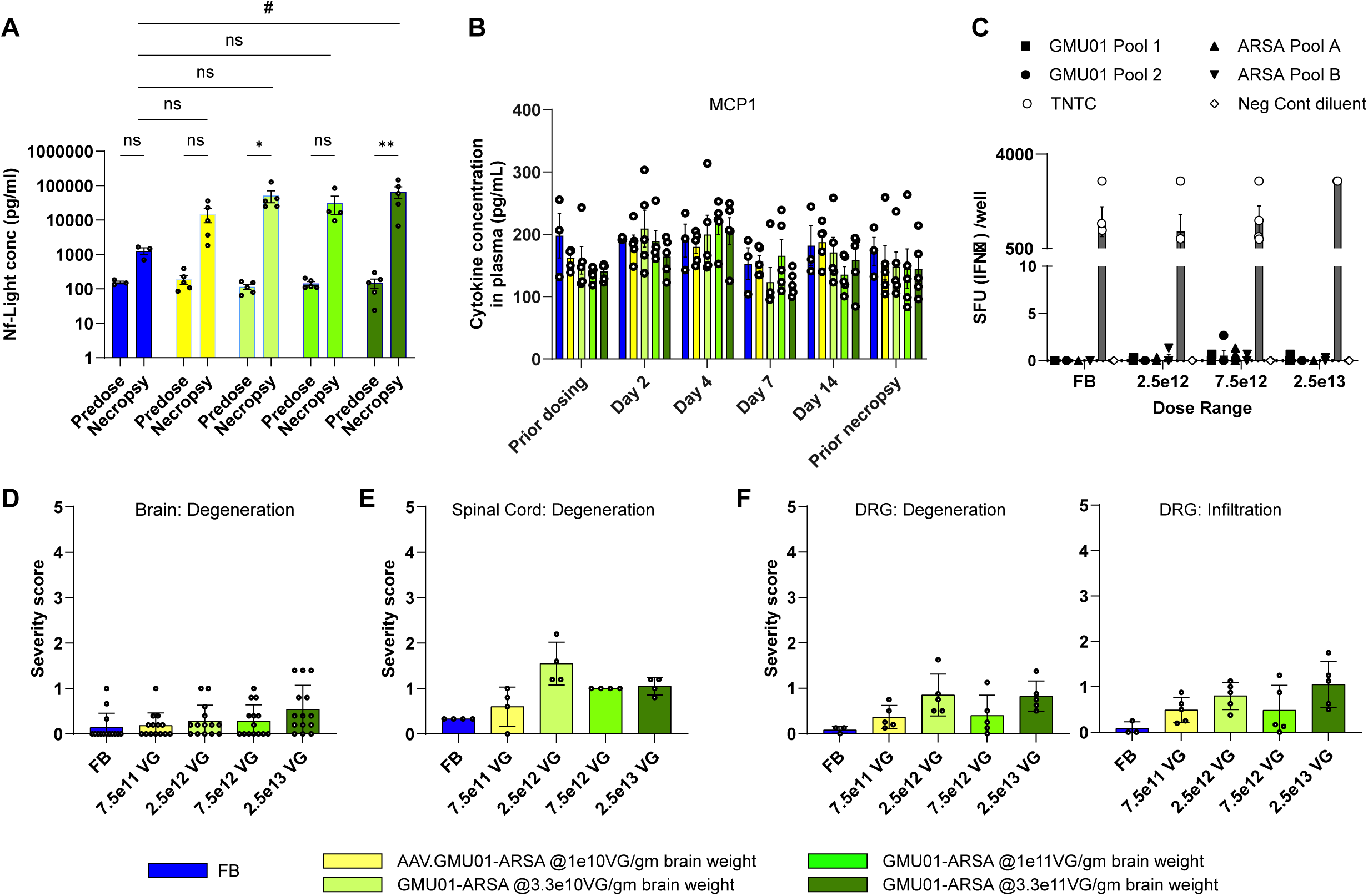
ICM infusion was well tolerated, resulted in expected Nf-L elevation, and did not trigger innate nor cell-mediated immune responses. (A) Pre-dose and at necropsy, CSF was collected and analyzed for Nf-L levels by Quanterix platform. Each data point represented Nf-L levels per animal, averaged across all NHPs in that group. (B) Plasma was isolated pre-dose, and at days 2, 4, 7, 14 post-dose, and at necropsy. The Luminex assay was used to determine the concentration of MCP1 (graphed) and IL-1b, IL-1RA, IL-6, IL-10, IL12/23 (p40), IL-15, IL-18, IFN-g, TNF-a, G-CSF, MCP-1, MIP-1b, GM-CSF, IL-2, IL-4, IL-5, IL-8, IL-13 and IL-17A (graphed in Supplementary Figure 13). Each data point represents cytokine concentration in that sample, averaged across all NHPs in that group. (C) PBMCs isolated from animals in the 1e11 and 3.3e11VG/gm brain weight dosing groups were subjected to IFN-γ ELISpot. (D-F) Histopathological findings in brain, spinal cord, and DRG; each data point represents the maximum severity of findings scored on 1-2 sections per animal. Severity scores refer to findings graded as 0= no findings, 1=minimal, 2=mild, 3=moderate, 4=marked, 5=severe. Error bars represent mean with standard error; Statistics: Two-way ANOVA with *Tukey’s and ^#^Dunnett’s multiple comparison test.

In conclusion, single administration of AAV.GMU01-*ARSA* by direct ICM injection results in widespread ARSA expression in CNS and is well tolerated in cynomolgus monkeys.

### AAV.GMU01-ARSA treatment results in potentially therapeutic levels of ARSA expression

Diminished ARSA function without clinical symptoms, referred to as ARSA pseudodeficiency, is present in approximately 1% to 2% of the global population (55, 56). Genetic and phenotypic comparisons indicate that individuals with ARSA enzyme levels between 5% and 20% of normal are largely asymptomatic (14, 57–62). To investigate the potential for AAV.GMU01-*ARSA* treatment to achieve therapeutic ARSA levels, we measured post-mortem ARSA protein in the brains (12 brain regions) of seven healthy human donors aged between 3 and 8 years old. Human brain regions from the 7 age-matched donors show an average of 1.9 fmol ARSA protein per 100ug total protein (**Figure 8A**). In comparison, brain-wide mean human ARSA protein levels in NHPs were 2.5 fmol (7.5e12VG/NHP) and 20.4 fmol (2.5e13VG/NHP) per 100ug total protein (**Figure 6E**). This corresponds to 1.2- and 8.1-fold higher ARSA protein levels than in the human samples (**Figure 8B**). Within-sample comparisons of human ARSA and endogenous cynomolgus (cyno)ARSA protein levels across 19 brain regions indicated that human ARSA protein levels were approximately 63% and 546% higher than the endogenous cynoARSA (**Figure 8C**) at the 7.5e12 and 2.5e13VG/NHP doses, respectively.

**Figure 8:**
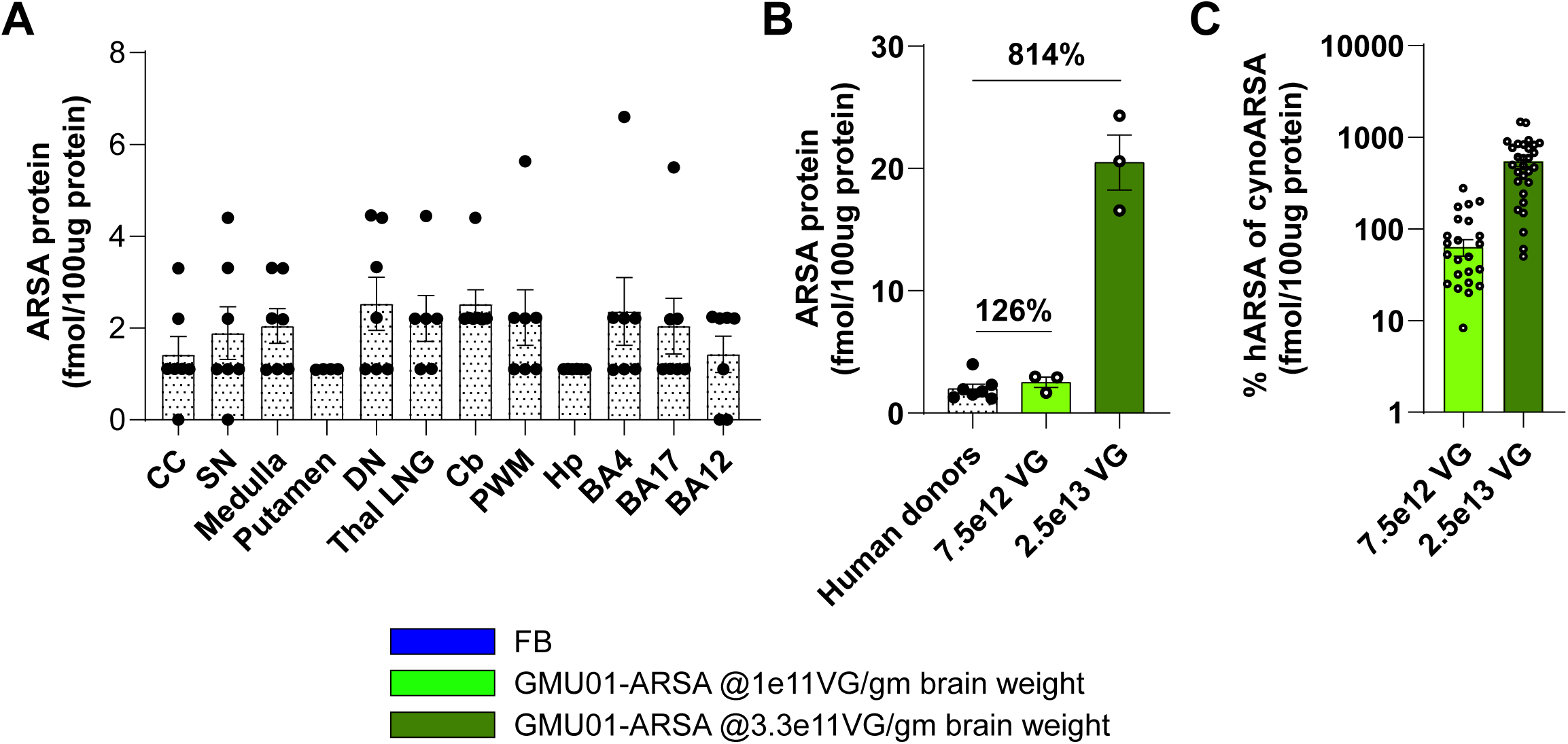
AAV.GMU01-*ARSA* effective doses are at potentially therapeutic levels. (**A**) LC-MS was performed to quantify human ARSA protein levels in 12 brain regions from 7 healthy 3- to-8 years old organ donors. Human tissue was received from the NIH NeuroBioBank at the University of Miami and the Sepulveda Research Corporation. (**B**) LC-MS was performed to quantify human ARSA protein levels in 30 tissue biopsy punches collected from brain (grey matter) of NHPs in dose-ranging study. Mean ARSA protein in each group is presented. (**C**) LC-MS was performed to quantify both human and endogenous *Macaca fascicularis* “cynoARSA” protein levels. Each data point is represented as a ratio of human ARSA to cynomolgus “cynoARSA” protein in that tissue punch and averaged across three animals in that group. Error bars represent mean with standard error. CC: corpus callosum; SN: substantia nigra; DN: dentate nucleus; Thal LNG: lateral geniculate nucleus of the thalamus; Cb: cerebellum; PWM: periventricular white matter; Hp: hippocampus; BA: Brodmann area.

We conclude that AAV.GMU01-*ARSA* treatment in NHPs results in clinically promising ARSA protein expression in the central nervous system.

## DISCUSSION

Adeno-associated viral vector gene therapy represents a promising approach for treating genetic diseases due to its ability to deliver sustained transgene expression, particularly within the CNS. Early evidence supporting the potential efficacy of ARSA replacement therapy in MLD emerged from studies in mouse models, where high doses of intravenously administered recombinant ARSA were shown to decrease sulfatide levels (63). However, enzyme replacement therapy (ERT) has demonstrated limited success in patients, primarily due to its inability to effectively cross the blood-brain barrier (64). Lentiviral hematopoietic stem-cell gene therapy (HSC-GT) has shown considerable efficacy in early-stage MLD, offering a continuous source of ARSA production. However, this approach is associated with significant risks stemming from the myeloablative conditioning required for transplantation (22). In contrast, AAV-mediated ARSA replacement therapy addresses these limitations by providing a safer, minimally invasive alternative. Preclinical studies across cellular, small animal, and NHP models have demonstrated the efficacy and safety of AAV-mediated ARSA delivery (63, 65–69).

The ability of AAV vectors to facilitate stable and sustained ARSA expression from a single administration promises long-term therapeutic benefits without the need for repeated treatments. This approach eliminates the peak-to-trough fluctuations in enzyme activity observed with ERT while circumventing the procedural risks associated with HSC-GT. Consequently, AAV-mediated ARSA replacement offers a robust and practical strategy for addressing the neurological and systemic manifestations of MLD.

### Advantages of AAV.GMU01 for ARSA Delivery

AAV.rh10 mediated gene replacement alleviated many long-term MLD-associated phenotypes in a mouse-model of MLD (33–35), but failed to demonstrate efficacy in the clinic despite long-lasting restoration of ARSA activity in the CSF (NCT01801709, (70)). This failure has been attributed to the inability of intracerebral delivery to achieve sufficient and widespread ARSA expression across the brain regions impacted by MLD (71). While well-tolerated in NHPs, the practicality and tolerability of multiple intracerebral injections in patients with extensive white-matter disease remain uncertain.

To overcome these limitations, AAV.GMU01 was specifically selected for its enhanced brain penetration, broad CNS biodistribution and robust transgene expression in NHPs following a minimally invasive ICM delivery. This approach enables targeted expression of ARSA in affected brain regions, achieving near-normal levels of ARSA activity without the need for direct intracerebral administration, which may be poorly tolerated in pediatric MLD patients. The clinical feasibility of ICM delivery is further supported by its use in ongoing trials for similar CNS conditions, including Phase 3 trials for GM1 gangliosidosis (NCT04273269) and MPS Type II (NCT03566043) in young children. Early-phase trials in adults with CNS disorders such as frontotemporal dementia (NCT04747431, NCT04408625), Parkinson’s disease (NCT04127578), MPS Type I (NCT03580083), and Gaucher Type II (NCT04411654) also demonstrate its broad applicability.

Importantly, intrathecal administration of AAV.GMU01 in NHPs resulted in lower vector exposure across the brain compared to AAV.rh10, while achieving superior biodistribution and transgene expression. This enhanced efficiency supports the potential for dose reduction, mitigating AAV-associated toxicity without compromising therapeutic efficacy. These attributes position AAV.GMU01 as a promising capsid for addressing the limitations of previous approaches and advancing ARSA gene replacement therapy for MLD.

### Preclinical Validation in Mouse and NHP Models

To evaluate the therapeutic potential of AAV.GMU01-*ARSA*, we conducted comprehensive pharmacology and efficacy studies in both mouse and NHP models. In mice, four studies were designed to model therapeutic intervention at various stages of neuropathological progression, reflecting pre-symptomatic and early-symptomatic MLD patients. Pharmacology studies in neonatal mice corresponded to the developmental age of the youngest intended patient population.

In the absence of ARSA activity, *Arsa* KO mice exhibit progressive sulfatide accumulation in the CNS and visceral organs, gliosis, microglial activation, and histopathological abnormalities, including degeneration and vacuolation in brain parenchyma. Auditory dysfunction, a hallmark of MLD, is also present in these models. Treatment with AAV.GMU01-*ARSA* restored ARSA activity to wild-type levels in various tissues, normalizing sulfatide levels, reducing neuroinflammatory markers, and reversing hearing impairment. Longitudinal studies demonstrated consistent ARSA-mediated sulfatase activity and sustained sulfatide reduction in the brain, spinal cord, CSF, and plasma. These findings support the applicability of CSF and plasma biomarkers for monitoring therapeutic efficacy.

Dose-ranging studies revealed robust increases in sulfatase activity and reductions in sulfatide deposits in neonatal and early-neuronopathic mice, supporting the therapeutic potential of this approach and informing dose selection for NHP studies. Notably, cross-correction extended the therapeutic reach of ARSA, with over 84% of the brain showing evidence of corrected cells, despite fewer than 10% being directly transduced. This process, mediated by uptake of extracellular enzyme via the cation-independent mannose-6-phosphate receptor (M6PR) (50–52), underscores the broad applicability of AAV.GMU01-*ARSA*.

Although bulk tissue assessments do not provide cell-specific resolution, our data suggest that AAV.GMU01 facilitates more productive transduction events compared to other capsids. This is consistent with prior studies, such as those by Goertsen et al., which demonstrated AAV.CAP-B10 achieved higher expression per cell despite lower cell transduction than AAV-PHP.eB, and where AAV.CAP-B22 outperformed AAV.CAP-B10 in brain expression despite similar DNA exposure (20). Kondratov et al. reported discrepancies between vector genome exposure and mRNA expression in a 29-capsid AAV library for the NHP CNS, showing that AAV.2i8 achieved higher expression than AAV5 at lower vector exposure (72), highlighting capsid efficiency as a key factor in improving therapeutic outcomes while mitigating AAV-associated toxicity.

In NHP studies, AAV.GMU01-*ARSA* was delivered via ICM infusion at varying doses. Dose-dependent increases in vector biodistribution and human ARSA mRNA and protein levels were observed throughout the CNS. Remarkably, human ARSA protein levels in NHP brains exceeded 126% of endogenous levels measured in healthy human donors at 1e11 VG/g brain weight, and over 800% at 3.3e11 VG/g brain weight. Such levels surpass those associated with ARSA pseudodeficiency, where individuals remain asymptomatic despite significantly reduced enzyme activity (14, 55–60). These findings suggest that even modest restoration of ARSA activity could have profound therapeutic effects in MLD patients, enabling effective treatment at lower doses.

The therapeutic potential of AAV.GMU01-*ARSA* was reinforced by its tolerability in NHPs. Although neuronal degeneration and mononuclear cell infiltrates were observed in some tissues, they were not associated with functional or behavioral deficits. Increased CSF Nf-L levels were detected post-treatment but were not dose-dependent, and no significant changes in plasma cytokine levels or cell-mediated immune responses were observed. These results suggest a favorable safety profile for AAV.GMU01-ARSA.

### Challenges and Limitations

While these preclinical studies highlight the promise of AAV.GMU01-*ARSA*, several areas warrant further investigation. The lack of cell-specific resolution in transgene expression assessments underscore the need for future detailed studies on cellular tropism. Moreover, GLP toxicology studies using clinically representative materials are essential for advancing this therapy to human trials.

Gene therapy inherently results in a mosaic distribution, with only a subset of transduced cells producing wild-type or elevated levels of ARSA to compensate for non-transduced cells. This mosaicism highlights the challenge of replicating natural low-activity states, such as those seen in individuals with ARSA pseudodeficiency, through therapeutic interventions. This limitation likely contributes to the incomplete normalization of sulfatide metabolites observed in this study, despite restored ARSA activity. Additionally, differences in sulfatide metabolism dynamics between mice and humans may contribute to incomplete sulfatide clearance in preclinical models, even when sufficient enzyme activity is present.

Cross-correction, where ARSA is taken up by non-transduced cells, is a key mechanism of therapy but may vary in efficiency across cell types, necessitating broader transduction or higher enzyme activity to achieve complete correction. Moreover, disease kinetics and specific metabolic demands in mouse models may not fully replicate human pathology. For example, murine ARSA exhibits approximately 3.5 times higher activity than human ARSA (73), potentially influencing the therapeutic response. Understanding the interplay between enzyme distribution, activity thresholds, and tissue-specific requirements will be essential for optimizing future therapeutic strategies.

Neurofilament light chain (Nf-L) serves as a sensitive biomarker for neuronal injury, neurodegeneration, and therapeutic efficacy in both preclinical and clinical settings. However, its interpretation in the context of AAV-based therapies is complex. Nf-L levels reflect neuronal damage or repair, which may lag behind metabolic changes, making it a delayed biomarker of disease progression or treatment response. In this study, Nf-L levels increased in both mouse and NHP models following AAV delivery, a phenomenon well-documented in the field. It is important to note that AAV administration is often associated with transient Nf-L elevations, confounding its use as a biomarker of disease repair during the early post-treatment phase. For example, in NHPs treated with AVB-101 (Aviadobio), Nf-L levels increased after intrathalamic AAV delivery but returned to baseline in serum and approached baseline in CSF after six months. Similarly, in clinical trials of AMT-130 (uniQure), patients experienced Nf-L elevations post-treatment that returned to near-baseline levels within 12–24 months, indicating a transient neuronal response to therapy. These observations highlight the need to interpret early Nf-L changes cautiously and to focus on long-term trends to assess therapeutic efficacy and safety accurately.

### Immune Considerations

Human ARSA shares 96% similarity with Cynomolgus monkey ARSA, raising the possibility of cross-reactive immune responses even in wild-type animals. While this high level of similarity suggests that central tolerance could mitigate immunogenicity, it does not entirely eliminate the risk of an immune response to the human ARSA transgene. This highlights the complexity of interpreting immune responses across species and underscores the need for immune monitoring during preclinical and clinical development.

In addition, the cross-reactive immunologic material (CRIM) status is a critical factor in shaping treatment strategies for MLD. CRIM-negative individuals, who lack endogenous ARSA protein expression, may be at higher risk of developing an immune response to the ARSA transgene. However, comprehensive data on CRIM status in MLD patients is limited. This gap stems partly from the challenges of measuring ARSA protein levels and the reliance on standardized dried blood spot (DBS) sulfatide and ARSA enzyme assays in newborn screening, which primarily aim to identify individuals at high risk of developing MLD rather than evaluate CRIM status. Existing data suggest that CRIM-negative patients may represent a small subset of the MLD population. A newborn screening study in Minnesota analyzing 100,000 blood spots identified 73 screen-positive samples, including 51 with pseudodeficiency variants (reduced but detectable ARSA protein), 20 heterozygous for pathogenic or unknown variants (reduced but detectable ARSA protein), and 2 homozygous for potentially pathogenic mutations (likely CRIM-negative)(74). This suggests that approximately 2.73% of newborns at high risk of MLD lack ARSA protein. Similarly, among European MLD patients, three common ARSA mutations were identified, with only one allele (allele I) associated with immunologically undetectable ARSA. Approximately 6% of MLD patients are homozygous for this allele, suggesting that 94% of MLD patients retain immunologically detectable ARSA (75, 76). These findings indicate that CRIM-negative individuals likely represent a small proportion of the overall MLD population, which has implications for clinical adoption of gene-replacement strategies like the one described in this study.

### Summary

This study highlights the therapeutic potential of AAV.GMU01-*ARSA* gene replacement therapy for MLD. Preclinical findings demonstrate that AAV.GMU01-*ARSA* effectively addresses key MLD-associated phenotypes, including sulfatide level normalization, reduction of neuroinflammation, and restoration of functional deficits, while maintaining a favorable safety profile in both mouse and NHP models. Notably, a single administration of AAV.GMU01-*ARSA* via ICM delivery in NHPs was well tolerated, achieving human ARSA expression at potentially therapeutic levels. These results provide a robust foundation for progressing to large-animal toxicology assessments with clinically representative materials. However, translating these promising preclinical outcomes to clinical applications requires careful consideration. Variations in enzymatic activity, biodistribution, and disease progression between species highlight the need for rigorously designed clinical trials to validate the safety and efficacy of this therapy in patients. In conclusion, AAV.GMU01-ARSA offers significant promise as a long-term, minimally invasive treatment for MLD, with the potential to transform patient outcomes and improve the lives of affected individuals and their families.

## METHODS

### Vector design

1. CBA-ARSA-WPRE-bGH: A 4397 nucleotide payload comprised of AAV2 Inverted Terminal Repeats (ITRs), a 1.7kb ubiquitous CBA promoter (CMV enhancer, chicken β-actin promoter, and a chicken β-actin/rabbit β-globin hybrid intron) expressing a functional copy of the human ARSA gene (ARSA), and having the bGH polyA and termination signal. The selection of the CBA promoter enables ARSA expression in most transduced cell types, ensuring system-wide expression. The transgene also incorporates an engineered Woodchuck Hepatitis Virus Posttranscriptional Regulatory Element (WPRE) to enhance ARSA expression in transduced cells.
2. CBA-Nanoluciferase-Flag-T2A-NLS-mCherry-bGH: A 4537 nucleotide payload comprised of AAV2 ITRs, a 1.7kb CBA promoter expressing both a FLAG-tagged Nanoluciferase (Nluc) and mCherry separated by the T2A ribosomal skipping peptide from a common ORF sequence with the bGH polyA and termination signal.
3. CBA-eGFP-bGH: A 4380 nucleotide payload comprised of AAV2 ITRs, the CBA promoter expressing a eGFP ORF and having the bGH polyA and termination signal. In vivo studies were performed utilizing material manufactured by transient transfection in adherent HEK293 cells.

#### Animal care and use

All experiments were conducted in AAALAC accredited institutions. Animals were handled in accordance with the rules and regulations of the IACUC, in compliance with the Animal Welfare Act, and adhered to principles stated in the Guide for the Care and Use of Laboratory Animals. All effort to minimize pain and distress were ensured in these purpose-bred animals. Only purpose-bred naïve cynomolgus NHPs were used in studies. NHPs were prescreened for AAV neutralizing antibodies, and seronegative animals were selected for the studies.

#### ARSA knockout mouse model

The model used mirrors the Gieselmann model (48) and was commissioned by Sanofi through Regeneron in 2012. The model is listed (Jackson Labs) as B6N.129P2(CBA)-^Arsatm1Gie/^J (*ARSA*^-/-^) and screened by “Loss of Native Allele (LOA) assay (77). The model was created through a disruption of *Arsa* Exon 4 near HindIII site (Genomic coordinates NCBIM37: chr15:89,302,907-89,307,855) with a loxP-hUBp-em7-Neo-polyA-loxP cassette in C57BL6 embryonic stem cell line, resulting in a positive clone (B-E9) with a Deletion of 44 bp for genotyping purposes NCBIM37:chr15:89,305,085-89,305,128. Disruption of the *Arsa* gene results in the progressive accumulation of sulfatide species in visceral organs, and the central nervous system, similar to the Gieselmann model (12, 78, 79). Gliosis and microglial activation is observed, with a notable auditory phenotype in line with the loss of neurons of the ventral cochlear nucleus, posterior part, and spiral ganglion, as seen in the Gieselmann model (12, 80).

### Rodent studies

1. *Single-dose studies*: Two-month old (pre-neuronopathic) *Arsa* KO mice underwent bilateral intracerebroventricular (ICV) injections (4 uL per hemisphere) of AAV.GMU01-*ARSA* at a dose of 1.6e11 VG per mouse (3.3e11 VG/gm brain weight). Six month-old (early-neuronopathic) *Arsa* KO mice underwent bilateral intracerebroventricular (ICV) injections (4 uL per hemisphere) of AAV.GMU01-*ARSA* at a dose of 5e10 VG per mouse (1e11 VG/gm brain weight). Control groups, including age-matched wild-type (WT) and Arsa KO mice, received doses of formulation buffer.
2. *Dose-ranging studies*: Neonatal (3 uL per hemisphere) or six-month old (5 uL per hemisphere) *Arsa* KO mice underwent bilateral intracerebroventricular (ICV) injections of AAV.GMU01-*ARSA* at a dose of 1e10, 3.3e10, 1e11 and 3.3e11VG/gram brain weight.
3. *Cross-correction study:* Thirteen month-old *Arsa* KO mice underwent bilateral intracerebroventricular (ICV) injections (5 uL per hemisphere) of AAV.rh10-*ARSA* at a dose of 1.6e11 VG per mouse (3.3e11 VG/gm brain weight).

### AAV administration in NHPs

1. AAV.GMU01 vs. AAV.rh10 comparison study: Cynomolgus monkeys (male, Mauritian 2-3 yr old, 2-3kg, n=3 per capsid) were implanted with an intrathecal catheter inserted at lumbar site and advanced to cervical 1-2 level for dose administration and/or CSF collection. The animals received two 2.5 mL injections at 0.125 mL/minute rate in Trendelenburg position through the ported lumbar catheter with an approximate 6-hour interval between injections.
2. Intra-CSF ROA optimization study: Cynomolgus monkeys (male, Vietnam, 2-3 yr old, 2-3kg, n=3 per route) animals were dosed by either bilateral intracerebroventricular injection (ICV) or direct cisterna magna injection (ICM). For ICV injections, animals were anesthetized and placed in the integrated head fixation frame with the ClearPoint (ClearPoint Neuro) array base placed on the cranium. The targeted dosing regions and cannula trajectory were determined using MRI scans. Prior to surgery, approximately 1 mL of CSF was removed. Bilateral ICV injections were perfumed sequentially under MRI guidance using SmartFlow NGS-NC-05 cannula (ClearPoint Neuro). A total of 1 mL of AAV.GMU01-NLuc-mCherry was administered at 0.125 mL/min flow rate into each lateral ventricle. For direct cisterna magna (ICM) injection, animals were placed in Trendelenburg position throughout dosing and for up to 15 minutes following dosing completion. Prior to dosing, approximately 1 mL of CSF was removed. A 24-gauge needle was inserted into the cisterna magna under fluoroscopy guidance. Animals were infused with a total of 2.0 mL AAV.GMU01-NLuc-mCherry at 0.125 ml/min flow rate while under anesthesia; A flush volume of 0.250 mL formulation buffer was given at the end of the dosing and the needle was left in place for at least 1-3 minutes before removal. Following the procedure, veterinary care was given for recovery.
3. Dose range finding study: Cynomolgus monkeys (3 males/2 females per group, Cambodia, 2-3 yr old, 2-3kg) were administered with AAV.GMU01-*ARSA* at 4 dose levels: 2.5e13 VG, 7.5e12VG, 2.5e12VG, 7.5e11VG via direct ICM as described above. Specifically, Animals were placed in Trendelenburg position during dosing. The dosing paradigm involved a single 2.5 mL infusion of AAV.GMU01-*ARSA* at 0.125 mL/min, followed by a 250 uL flush with formulation buffer.

#### NHP sample collection

At necropsy, animals were perfused with chilled PBS pH 7.4, and their brains were cut in a brain matrix into 4 mm coronal slices and hemisected. The left hemispheres were frozen on dry ice for biochemical analysis, while the right hemispheres were fixed in 10% neutral buffered formalin for 36-48 hours at room temperature before embedding in paraffin blocks for histopathology and immunohistochemistry analysis. The spinal cord and DRGs from 4 levels (cervical, upper thoracic, lower thoracic and lumbar), as well as peripheral tissues were collected as frozen and fixed samples. Tissue punches were collected using 3 mm biopsy punches from indicated brain regions on the frozen brain slices, as well as from 4 levels of spinal cords, DRGs and peripheral tissues. Paraffin tissue sections were stained for hematoxylin and eosin and submitted for histopathology analysis.

#### NHP Histopathology

Pathology evaluations were conducted by a board-certified veterinary pathologist at HistoWiz (Long Island City, NY) or StageBio (Mt Jackson, VA) on hematoxylin and eosin-stained slides. For each animal, eight key brain regions (cerebral cortex, basal ganglia, thalamus, hippocampus, midbrain/pons, cerebellum, medulla and major white matter tracts), as well as spinal cord and DRGs from cervical, upper thoracic, lower thoracic and lumbar levels were examined for axon/neuron degeneration and/or inflammation/cell infiltration. A minimum of 8 DRGs (2 from each level) were evaluated. Microscopic findings were graded as 0 for absence of lesion, 1 for minimal, 2 for mild, 3 for moderate, 4 for marked, and 5 for severe.

#### NHP GFP Immunohistochemistry (IHC)

For GFP IHC, antigen retrieval was performed on FFPE slides from brain using EDTA solution (pH.9.0) for 20 minutes at 90°C. After blocking with 3% hydrogen peroxide for 10 minutes and 5% horse serum for 45 minutes, the slides were incubated with GFP antibody (ThermoFisher A-11122) at 1:500 dilution for one hour at room temperature, followed by incubation with anti-rabbit HRP (Abcam ab6721) at 1:200 for one hour at room temperature. The color development for GFP signal was achieved by incubating slides in DAB solution (Thermofisher 34002) for 3 minutes at room temperature. Brightfield images were captured by an Aperio AT2 image scanner.

#### Tissue homogenization

Tissues were homogenized at 4C in cold TE buffer (10mM Tris pH7.4, 1mM EDTA) in 2mL tubes containing 1.4 mm ceramic beads using an Omni Beadruptor set for 20 second cycles 4.7 oscillation/sec. Following homogenization, aliquots were frozen at –80C until use.

#### Tissue solubilization for Sulfatase Activity

To homogenized tissue, Nonidet P-40 was added to 0.1% final concentration, allowed to solubilize at 4C for 1.5 hours on an orbital shaker, then centrifuged at 18,000 *x g* for 20 minutes. Supernatant was removed and transferred to fresh Eppendorf tube on ice prior to BCA and Sulfatase activity assays.

#### BCA Assay

Total protein concentration determined by BCA (bicinchoninic acid) assay (Thermo Scientific 23227) using 10ul supernatant diluted in water. Colorimetric detection of the cuprous cation (Cu^1+^) by bicinchoninic acid (BCA) by absorbance at 562 nm. Molecular Devices SpectraMax 340PC-384 with SoftMax Pro version 5.4.4 software used to read 96 well microtiter plate.

#### Nanoluciferase activity

Crude tissue homogenates were assayed for nanoluciferase activity using the Nano-glo luciferase assay system (Promega) according to manufacturer’s instructions. Luminescence was read using the Cytation C10 and was normalized to total protein content as measured using the BCA assay.

#### Sulfatase Activity

Ten microliters of clarified supernatant was assayed for total sulfatase activity using sulfatase activity assay for hydrolyzed 4-Nitrocatechol (PNC) from 4-Nitrocatechol Sulfate (PNCS) substrate. (Abcam Ab204731). Activity determined by hydrolyzed 4-Nitrocatechol (PNC) of sample relative to PNC standard curve and read absorbance at 515 nm. Sulfatase activity reported as mU/mg (nmol/min/mg). Molecular Devices SpectraMax 340PC-384 with SoftMax Pro version 5.4.4 software used to read 96 well microtiter plate.

#### Lipid extraction

An aliquot of tissue homogenate (or fluid) was extracted with 20-100X extraction solution (5 mM Ammonium Formate, 0.2% Formic Acid in Acetonitrile:Methanol (70:30), supplemented with 10 ng/ml C17-sulfatide). After mixing vigorously for 10min, samples were sit for 5min and vortexed again quickly (30s). All samples were centrifuged at 8,400 rpm for 10 min at 4°C. An aliquot of the supernatant (200uL) was transferred into a deactivated Q-sert Vial for LC-MS analysis. Standard curve (linearity range 0.03-1000 ng/mL, 2x-serial dilutions) were prepared in the same extraction solution using sulfatides with C16, C18, C24 and C24:1chain length (Matreya) and C18 2R-OH sulfatide and lyso-sulfatide (Avanti).

#### LC-MS analysis of Sulfatides

A Waters Acquity UPLC system (Milford, MA) was coupled to a Qtrap 6500 mass spectrometer system (Framingham, MA) equipped with a ESI source operated in negative ion mode with the following parameters: curtain gas 25.0; ionSpray voltage −4.5 kV; temperature 500°C; ion source gas: 50 and 70; declustering potential −80V; entrance potential - 10V; collision energy −155V; and collision cell exit potential −15V. The sulfatide species were separated on a Waters Acquity UPLC BEH Amide (1.7 µ, 2.1 × 100 mm, P/N: 186004801). The autosampler and column oven were maintained at 10°C and 20°C, respectively. The mobile phase consisted of Solvent A: 5 mM Ammonium Formate in 95:5 of Acetonitrile:Water, and Solvent B: 5 mM Ammonium Formate in 90:10 Methanol:Water. The gradient program started at 0% B and hold for 2min, followed a linear curve from 0%B to 100% B. After 1 more min at 100% B, it re-equilibrated at 0%B. All sulfatide species are normalized to the internal standard of C17-sulfatide, and calculated by the standard curve of the corresponding standard sulfatides.

#### GFP ELISA

Measurement of GFP protein was completed using the GFP SimpleStep ELISA® Kit from Abcam (ab171581). Samples homogenized in TE buffer as described above were diluted in complete cell extraction buffer as follows: grey matter 1:5, spinal cord 1:5, heart: 1:5, liver 1:20. Samples were read at 450nm on a SpectraMax plate reader (Molecular Devices) using Softmax software.

#### Genomic DNA (gDNA) isolation

DNA was isolated from 50ul tissue homogenate using QIAmp 96 DNA QIAcube HT kit (cat# 51331) on a Qiacube HT according to manufacturer’s protocol “QIAamp® 96 DNA QIAcube® HT Handbook”. DNA concentration then was measured via absorbance with NANODROP 8000 (Thermo Fisher Scientific).

#### Digital PCR

Vector genome copies were determined from extracted DNA by dPCR using the QIAcuity Eight digital PCR System (Qiagen, Inc.) gDNA was analyzed in a 12ul reaction volume. Probes for bGH (AAV) and an endogenous mouse or cynomologous reference gene (Integrated DNA Technologies, Inc.) were used to measure gene copy number and calculate vector genomes per cell (2X bGH copies divided by reference gene copies.

#### Total RNA isolation

An aliquot of TE tissue homogenate was used for RNA extraction from the QIAcube HT (Qiagen, Inc.) using the RNeasy 96 QIAcube HT kit (#74171) according to manufacturer’s protocol “RNeasy 96 Qiacube HT Handbook” with on-plate DNase digestion, with noted QIAzol (Qiagen #79306) chloroform RNA extraction according to the QIAgen “QIAzol Handbook” steps 1-7 preceding and transferring the supernatant to the S-block for QIAcube HT RNA purification.

#### RT-dPCR

Isolated total RNA copies were reverse transcribed to cDNA (Qiagen One Step Advanced Probe kit, Qiagen #250132) and quantitated for copies/ul using the QIAcuity Eight digital PCR System (Qiagen, Inc.) utilizing specific qPCR primer and probe sets for *hARSA* (Integrated DNA Technologies, Inc.), gene normalizer (*Hprt*, mouse or cynomologous specific), inflammatory markers (*Gfap, Aif1*), and marker of lysosomal health (*Lamp1*). Total reaction volume is 12ul per well. The 8.5k 96 well nanoplate (Qiagen #250021) was placed into QIAcuity instrument and RT-dPCR performed using manufacture suggested cycling. RNA copies for each target gene were analyzed automatically by the software. Target gene was then normalized to the housekeeping gene using the QIAcuity software suite.

#### Cross correction analysis

Adjacent 5 µm FFPE sections were processed for either ARSA IHC and DAPI or *ARSA* ISH (against WPRE mRNA) and DAPI. Individual tiles were then stitched to obtain sagittal sections for both IHC and ISH images. These images were then registered globally with a transformation matrix (rotational and translational parameters) to align the IHC and ISH images. Individual tiles within the sagittal sections were then registered locally using DAPI to ensure good alignment between individual IHC and ISH tiles. In each tile, IHC positive cells and ISH positive cells were estimated by thresholding parameters empirically determined across the entire sagittal sections. In the case ISH, IHC positive cells had to have at least 20% overlap with a nuclei to be counted as a bonafide ISH signal. For IHC images, negative controls with no ARSA staining were used to determine thresholding parameters. Cross-correction factor which is the ratio of IHC positive cells to ISH positive cells was estimated for each tile and represented as a heat map.

#### ARSA LC-MS

Tissue homogenates from humans and non-human primates (NHPs) were processed for proteomic analyses using a stepwise protocol involving reduction, alkylation, and overnight digestion with Trypsin/Lys C. Sample handling was performed with the MINI 96 (Integra Biosciences) digital pipettor to enable high-throughput analysis. Digested peptides, spiked with synthetic heavy peptides labeled at the C-terminal R/K (13C15N), were clarified using EvoTip trap columns. These were connected online to an Evo 8 cm analytical column and separated using a preset 21-minute analytical gradient (60SPD) on the Evosep One nanoLC system. Absolute quantification of human and NHP ARSA proteins was attempted through an in-house multiplex LC-MS assay, leveraging high-field asymmetric waveform ion mobility spectrometry (FAIMS)-based gas-phase ion separation in high-resolution mode. Target peptide elution was empirically determined via chromatographic retention time (RT), and FAIMS-separated gas-phase ions were analyzed through parallel reaction monitoring (PRM).

For FAIMS-PRM, compensation voltages (CVs) for unique human and NHP peptides were optimized by ramping DC voltages from −50 CV to −30 CV to identify the optimal values for each peptide. While human brain homogenates followed a similar process, FAIMS was not utilized for their analysis. MS1 data acquisition for all samples occurred at high resolution (240K), with tMS2 data acquired at 30K resolution. The accuracy and precision of the FAIMS-PRM assays were evaluated using a 12-point calibration curve with linear ranges of 0.02–50 fmol/µL for the human-unique peptide (QSLFFYPSYPDEVR) and 0.1–25 fmol/µL for the NHP-unique peptide (GGLPLEEVTLAEVLAAR). Quality controls (QCs) at low, medium, and high levels were prepared by spiking surrogate peptides into a pooled matrix. The ratio of endogenous to surrogate peptides was obtained from Skyline, with ARSA expression values calculated using a single-point calibration method:

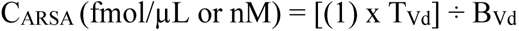

R= L/H; L= endogenous and H= surrogate (heavy)

S_O_= spike (H) on-column (fmol)

*Vd0*= digests volume (on-column)

T_Vd_ (µL) = total volume (µL) of starting materials

BVd (µL) = Total volume (µL) of brain homogenate (undigested) provided

To ensure precision and reproducibility, we achieved sub-nM quantification using a cutting-edge mass spectrometry platform, the Thermo Exploris 480 integrated with FAIMS and the Evosep nanoLC system. Isotope-labeled and unlabeled peptides for human-specific, NHP-specific, and shared peptide sequences were synthesized for calibration and QC. The human-unique peptide (QSLFFYPSYPDEVR) demonstrated a linear range of 0.02–50 fmol/µL, with a limit of quantification (LOQ) of 0.02 fmol/µL, while the NHP-unique peptide (GGLPLEEVTLAEVLAAR) exhibited a linear range of 0.1–25 fmol/µL and an LOQ of 0.1 fmol/µL. QC thresholds for high, medium, and low levels were defined as 5, 2.5, and 0.25 fmol/µL, respectively.

Sensitivity was validated through linear dynamic range and LOQ determination, with QC data consistently maintaining standard deviations within 10% for the human peptide and 17% for the NHP peptide, adhering to the FDA acceptability criteria of ±20%. These results highlight the assay’s reliability, enabling accurate quantification of peptides at sub-nM concentrations and reinforcing its clinical relevance.

### ELISpot procedure

- PBMC culture & immune stimulus: Frozen PBMCs were thawed at 37°C for 10 minutes and diluted in RMI-1640 medium containing 10% heat inactivated-FBS. Cells were plated at 100ul/well, in triplicate, to precoated 96 well ELISPOT plates purchased with anti-hIFN-gamma Single-Color ELISPOT kit (ImmunoSpot) for human (will cross react with NHP).
- Peptide Library & Positive Control Preparations: Peptide libaries for the AAV.GMU01 capsid, and the hARSA gene product were generated by Mimetope and consisted of 15mer peptides spaced every 3 amino acids of the sequence. Each individual peptide was prepared in 80% DMSO at 50mg/ml.
- Positive Control Preparations: Ionomycin calcium salt from Streptomyces conglobatus [Sigma Aldrich I3909] and PMA [Phorbol-12-myristate-13-acetate – Calbiochem #5005820001] was used as positive controls. Stocks were prepared in DMSO and diluted in RMI-1640 medium to 4uM Ionomycin/100nM PMA
- ImmunoSpot hIFN-g: Single-Color ELISPOT kit protocol was followed for staining and detection, as described by manufacturer. Plates were then imaged and counted on the C.T.L ImmunoSpot S6 Universal M2 analyzer with ImmunoSpot 7.0.28.4 Analyzer Proffesional DC; Immunospot 7.

#### Cytokine panel

Quantification of pro-inflammatory cytokines in plasma was performed by Charles River by Luminex Assay. The Analyte Specific Procedure describing the procedure to determine the concentration of IL-1β, IL-1RA, IL-6, IL-10, IL-12/23 (p40), IL-15, IL-18, IFN-γ, TNFα, G-CSF, MCP-1, MIP-1β, GM-CSF, IL-2, IL-4, IL-5, IL-8, IL-13 and IL-17A in Cynomolgus monkey plasma by Luminex assay can be provided upon request.

#### NfL assessment

NF-L protein level was measured using Simoa NF-Light*TM* V2 Advantage kit from Quanterix (Item# 104073). Mouse plasmas sample were diluted 20-fold in kit sample diluent. NHP CSF sample were diluted 160-fold in Lysate diluent C reagent purchased from Quanterix (item# 103360). Diluted samples were analyzed on Simao HD-X analyzer.

#### Immunofluorescent RNA/Protein integrated co-detection assay

Co-detection of WPRE RNA with payload proteins (human arylsulfatase A or mCherry) and cell type markers from rodent and non-human primate (NHP) brain FFPE slides was performed using a Leica BOND RX automated stainer (Leica Biosystems). The multiplexing protocol was programmed to streamline *in situ* analysis of transgene RNA and a continuous IHC staining process for the co-detection of RNA and protein markers. The RNAscope 2.5 LS multiplex fluorescent kit (ACD, 322800) and the Bond polymer refine detection kit (Leica Biosystems, DS9800) were utilized in this automated protocol. Specific WPRE probes (rodent: ACD, 450268; NHP: ACD, 410058) targeted the 3’ UTR of the transgene RNA during the *in situ* hybridization (ISH) step. Detection and visualization were achieved using TSA Vivid Fluorophore 647 (Biotechne, 75271KIT). Sequential IHC steps involved the use of antibodies against human ARSA (1:250, R&D Systems, AF2485) or mCherry (1:250, MAB131873), along with cell marker antibodies (1:500) for identifying neurons, astrocytes, microglia, and oligodendrocytes. DAPI was used for nuclear counterstaining. This integrated protocol enables precise co-detection of transgene RNA and protein with cell-specific markers, providing a comprehensive analysis of expression and cellular localization across rodent and NHP brain tissue samples.

| Kits | Vendor | Catalog No. |
| --- | --- | --- |
| RNAscope™ LS Multiplex Fluorescent | ACD | 322800 |
| Wpre-O1 probe | ACD | 450268 |
| Bond Polymer Refine Detection Kit | Leica Biosystem | DS9800 |
| BOND Research Detection System | Leica Biosystem | DS9455 |
| TSA Vivid 650 | R&D Systems | 75271KIT |
| RNAscope™ LS Multiplex Fluorescent | ACD | 322800 |
| Wpre probe | ACD | 410058 |

| Primary antibody |  |  |  |
| --- | --- | --- | --- |
| Targets | Host | Vendor | Catalog No. |
| Human arylsulfatase A/ARSA, pAb | Goat | R&D Systems | AF2485 |
| Anti-NeuN antibody [EPR12763], mAb | Rabbit | Abcam | ab177487 |
| Iba1/AIF-1 (E4O4W) XP, mAb | Rabbit | Cell Signaling Technology | 17198S |
| S100B (E7C3A) | Rabbit | Cell Signaling Technology | 42397 |
| Anti-mCherry [1C51] | Mouse | Millipore Sigma | MAB131873 |
| Anti-NeuN antibody [EPR12763], mAb | Rabbit | Abcam | ab177487 |
| Iba1/AIF-1 (E4O4W) XP, mAb | Rabbit | Cell Signaling Technology | 17198S |
| S100B (E7C3A) | Rabbit | Cell Signaling Technology | 42397 |
| Anti-Olig2 [EPR2673] | Rabbit | Abcam | ab109186 |
| Secondary antibody |  |  |  |
| Targets | Host | Vendor | Catalog No. |
| Anti-Rabbit IgG – Alexa Fluor 488 | DK | Thermo | AF32790 |
| Anti-Goat IgG – Alexa Fluor 555 | DK | Thermo | AF32816 |
| Anti-Mouse IgG – Alexa Fluor 555 | DK | Thermo | AF32773 |

#### Microscopic Imaging

Images were acquired using Zeiss Axio Scan. Z1 slide scanner with Colibri 7 fluorescence light source (Carl Zeiss MicroImaging). Images were acquired at 20x magnification and exported into ZEN Microscopy Software 3.3 (Carl Zeiss MicroImaging) for brightness/ contrast adjustment to increase the clarity and to reflect true rendering.

### Single-nuclei RNA sequencing data

- *Nuclei Isolation:* Tissue samples were stored at −80°C. For tissue lysis and washing of nuclei, sample sections were added to 1 mL Nuclei PURE lysis buffer (Sigma, NUC-201) and thawed on ice. Samples were then Dounce homogenized with PestleAx20 and PestleBx20 before transfer to a new tube, with the addition of additional lysis buffer. Following incubation on ice for 15 minutes, samples were then filtered using a 30 mM MACS strainer (Fisher, NC0642012), centrifuged at 500xg for 5 minutes at 4°C using a swinging bucket rotor (Sorvall Legend RT, Thermo Fisher), and then pellets were washed with an additional 1 mL cold lysis buffer and incubated on ice for an additional 5 minutes. A 1.8M sucrose buffer was created and 1.8ml was mixed with 1ml of cell suspension. 1ml of sucrose buffer was deposited to ultra centrifuge tube (Fisher, NC9157569) then the cell suspension was layered on top (2.8ml). The suspension was then centrifuged at 30,000xg (17,800 rpm) for 45min at 4C, Rotor SW55ti. Supernatant was then aspirated off, and pellet was resuspended in 1ml of wash buffer (Nuclei Pure storage buffer, Sigma NUC-201) and spun at 500xg for 5 min at 4°C. Supernatant was aspirated and 0.5ml of wash buffer was added for resuspension. For NeuN/Dapi and FACS sorting, from 0.6 mL nuclei sample, 540 mL, 30 mL, and 30 mL were aliquoted into tubes for sample and controls and then 10X Dapi/NeuN buffer was added to tubes for a final 1X concentration. Tubes were then incubated on ice for 30 minutes, with inversion every 10 min. Following incubation, samples were spun at 500xg for 5 min, supernatant removed, and samples were resuspended in 600 ul Wash buffer for samples (300 ul for control tubes). Nuclei then underwent filtering through 70um filter and filtered again through 40um filter (Fisher, 14100150) just before sorting using BD Bioscience InFlux Cell Sorter.
- *Sample QC and library Preparations*. Library preparation and NovaSeq Sequencing: Libraries were prepared according to 10xGenomics protocol for Chromium Single Cell 3’ Gene Expression V3.1 kit. NovaSeq sequencing was performed according to illumine NovaSeq 6000 protocol. UMI count matrices generated by Cellranger V5.0.0.
- *Single-nuclei RNA sequencing data preprocessing.* CellBridge^1^, was used to perform pre-processing of the data. CellBridge, in brief, performs QC thresholding (minimum UMI = 750, percent.mt < 20, nFeature_RNA<250), a log normalization of count data (LogNormalize function with a scaling factor of 1e+06), FindVariableFeatures (nfeatures=2000), RunUMAP (dims = 1:30, res=0.7, k=20), and Harmony (by sample) and outputs a R-based Seurat object. Automated celltype calling was performed by SARGENT^2^, a cell-by-cell geneset-based classification algorithm. Genesets for cell-type annotation were acquired from previously annotated cell types^3^ and filtered for the top positive (log2FC > 1.5, pct.1/pct.2 > 10 | pct.1-pct.2 > 0.5, slice_max(n=50) and negative markers (log2FC < - 1.5, pct.2/pct.1 > 10 | pct.2-pct.1 > 0.5). All code for cell type calling and processing to be provided in Sanofi-Public github.
- *Analysis Pipeline.* Primary data visualization pipelines in R were run under the R 4.3.2 environment. UMAPs were plotted using native Seurat (5.1.0). Calculating percent GFP expression was calculated using Percent_Expressing function in the scCustomize package and sumarrySE function in the Rmisc package. Cell type proportion comparisons were visualized using ggplot2 and significance comparisons were performed using an unpaired t-test for the cell types considered in the plot. For differential expression analyses between conditions, a custom wrapper function using edgeR (4.0.16)/limma (3.58.1) was created. In brief, this function performs pseudobulk aggregation using the function AggregateExpression on a Seurat object’s raw count data, per condition_cellstate. Then, we created a pairwise design and contrast matrix and the equation ∼0+comparsioncellstates with no input covariates into the analyses. A standard recommended pipeline for creating a negative binomial generalized linear model with F-test using filterByExpr, calNormFactors, estimateDisp, and glmQLFit was then called and individual glmQLFTests were run in parallel using foreach (1.5.2) and doParallel (1.0.17) packages.

#### NIH NeuroBioBank

Human tissue was received from the NIH NeuroBioBank at the University of Miami and the Sepulveda Research Corporation.

#### Statistical Tests

In the provided manuscript, various statistical tests were used across the figures to analyze the data and notated in the figure legends. We widely applied One- and Two-way ANOVA, either with Tukey’s, Sidak’s or Dunnett’s test, where appropriate. Correlation analysis was applied to assess the relationship between vector genome exposure and transgene expression or protein levels. All error bars in graphs represent mean with standard error.

Data and materials are available upon reasonable request. Contact.

## Author contributions

Conceptualization: SR, MG, CM

Methodology: SR, JA, JB, LG, DG, YC, SA, QT, COR, BZ, GG

Investigation: JA, JB, MAR, LG, DG, MH, YC, EW, SA, QT, RT, YL, JH, DDB, EC, JA, AR, SN, BN, DW, SCS, MT

Data Curation: JA, JB, DG, SA, QT, JH, JA, MT, SCS

Analysis: SR, JA, DG, SA, JH, JA, BE, MT, GG, SCS

Visualization: SR, JA, SA, JH, JA, BE, MT, SCS

Supervision: SR, MG, CM

Writing – original draft: SR, JA, JB

Writing – review & editing: SR, MG, BE, JH

## Acknowledgments

Human tissue was received from the NIH NeuroBioBank at the University of Miami and the Sepulveda Research Corporation. The following Sanofi employees provided support: peer review of pathology reports was supported by Karamjeet Pandher and Rachel Peters. Preclinical safety support was provided by Basel Assaf and Christopher Thompson. Dose selection support was provided by William McCarty. Statistics review was provided by Weiliang Qiu. Review and writing support were provided by Edith Pfister and Michael Fleming. Project administration support was provided by Kristin Radzwill. Portfolio supervision was provided by Evis Havari. Clinical support was provided by Nazem Atassi and Elena Gargaun. IP support was provided by Alejandro Martinez.

