## Supplementary Figures and Tables for "AAV-mediated ARSA replacement for the treatment of Metachromatic Leukodystrophy"

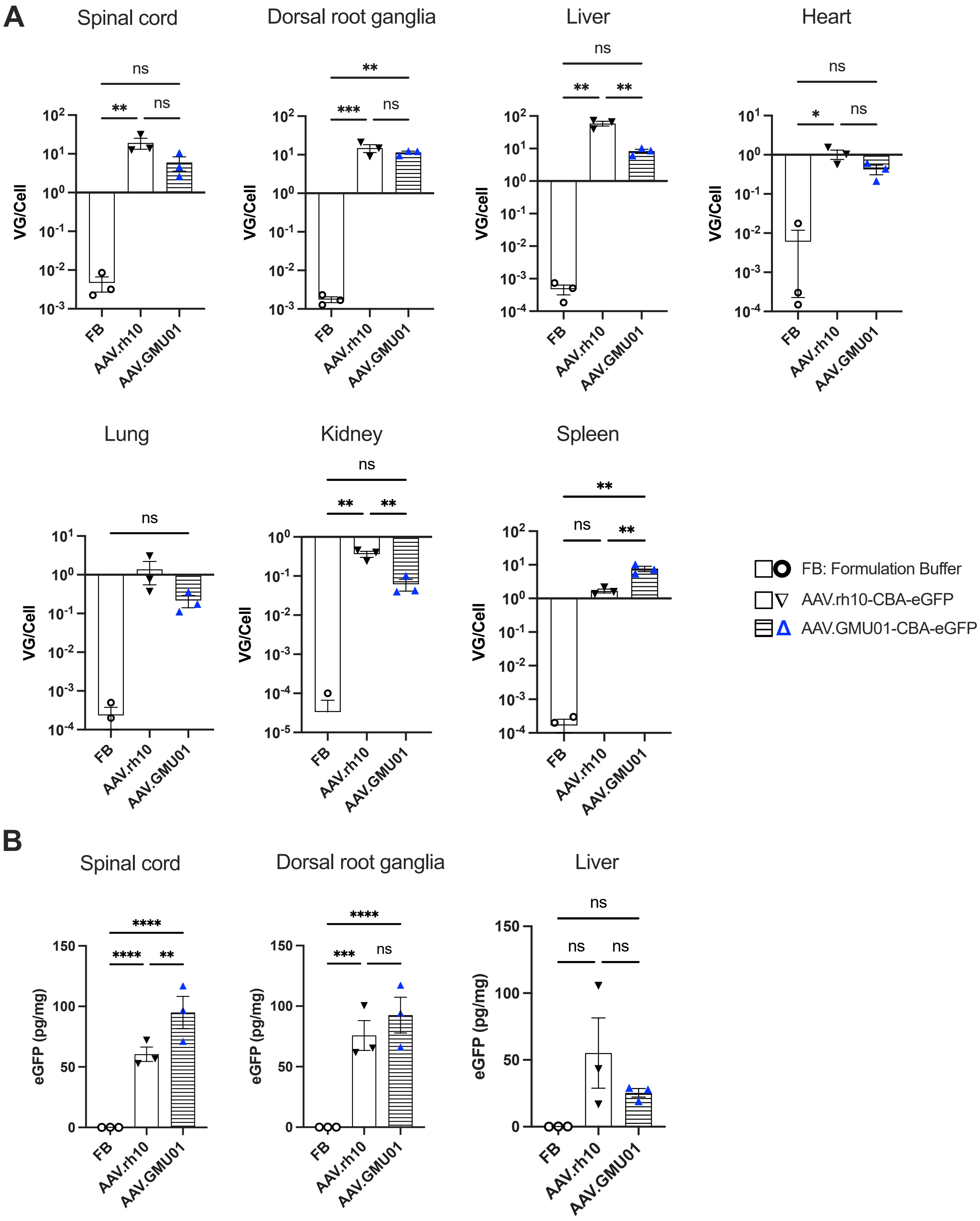
A
Spinal cord
Dorsal root ganglia
Liver
Heart
Lung
Kidney
Spleen
VG/Cell
VG/cell
ns
**
***
*
FB
AAV.rh10
AAV.GMU01
FB: Formulation Buffer
AAV.rh10-CBA-eGFP
AAV.GMU01-CBA-eGFP
B
Spinal cord
Dorsal root ganglia
Liver
eGFP (pg/mg)
****
**
***
ns
150
100
50
0

**Supplemental Figure 1. The novel capsid AAV.GMU01 shows higher transgene expression in the spinal cord and DRG of NHPs, compared to AAV.rh10.** Cynomolgus monkeys (Male, Mauritian 2-3yr old, 2-3kg) seronegative for AAV.rh10 and AAV.GMU01 were dosed intra-thecal at the cervical level 1-2 junction using a ported intrathecal catheter inserted at the lumbar region. Animals were dosed in the Trendelenburg position. One dose of AAV.GMU01-CBA-eGFP, AAV.rh10-CBA-eGFP or formulation buffer was administered at 2.75e13VG/NHP (3.65e11VG/gm brain weight). Two weeks post-dosing, animals were euthanized, and samples were assessed for vector genome exposure and eGFP expression. Spinal cord and DRG from 7 segments along the spinal column. (**A**) AAV vector exposure by dPCR - (**B**) eGFP expression by ELISA. Error bars represent mean with standard error. Two-way ANOVA with Tukey's multiple comparison test. *$p$<0.05; **$p$<0.01; ***$p$<0.001; ****$p$<0.0001.

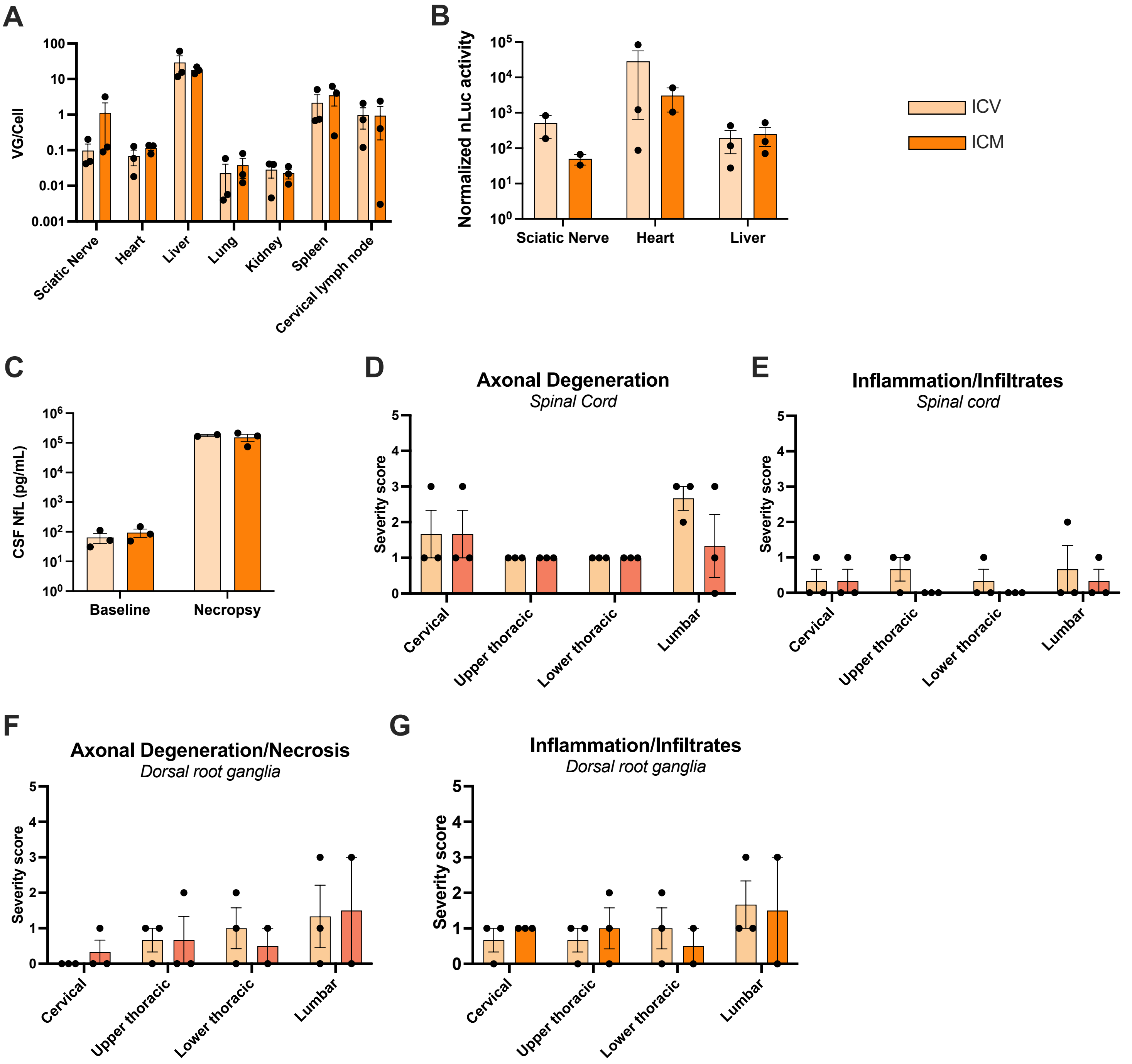

A
VG/Cell
100
10
1
0.1
0.01
0.001
Sciatic Nerve
Heart
Liver
Lung
Kidney
Spleen
Cervical lymph node
B
Normalized nLuc activity
Sciatic Nerve
Heart
Liver
ICV
ICM
C
CSF NfL (pg/mL)
Baseline
Necropsy
D
Axonal Degeneration
Spinal Cord
Severity score
Cervical
Upper thoracic
Lower thoracic
Lumbar
E
Inflammation/Infiltrates
Spinal cord
Severity score
Cervical
Upper thoracic
Lower thoracic
Lumbar
F
Axonal Degeneration/Necrosis
Dorsal root ganglia
Severity score
Cervical
Upper thoracic
Lower thoracic
Lumbar
G
Inflammation/Infiltrates
Dorsal root ganglia
Severity score
Cervical
Upper thoracic
Lower thoracic
Lumbar

**Supplemental Figure 2: AAV.GMU01 shows widespread vector biodistribution and transgene activity in peripheral tissues.** Cynomolgus monkeys (Male, Vietnam, 2-3yr old, 2-3kg) seronegative for AAV.GMU01 were dosed with AAV.GMU01-CBA-NLuc-mCherry at 2.0e13VG/NHP (2.75e11 VG/gm brain weight) either by bilateral intracerebroventricular injection (ICV) or direct injection to the cisterna magna (ICM). Four weeks post-dosing, animals were euthanized, and tissues were flash-frozen from peripheral organs. (**A**) AAV vector exposure by bGH-dPCR, (**B**) nanoluciferease (nLuc) activity, normalized to total protein measured by bicinchoninic acid (BCA) assay. (**C**) Cerebrospinal fluid (CSF) samples were collected both prior to dosing ("prestudy") and at the four-week necropsy timepoint and were assayed by Quanterix Simoa to quantify neurofilament light chain (NfL), a measure of neuronal injury. (**D-G**) Histopathological findings in brain, spinal cord, and DRG; each data point represents the maximum severity of findings scored on 1-2 sections per animal. Severity scores refer to findings graded as 0= no findings, 1=minimal, 2=mild, 3=moderate, 4=marked, 5=severe. Error bars represent mean with standard error.

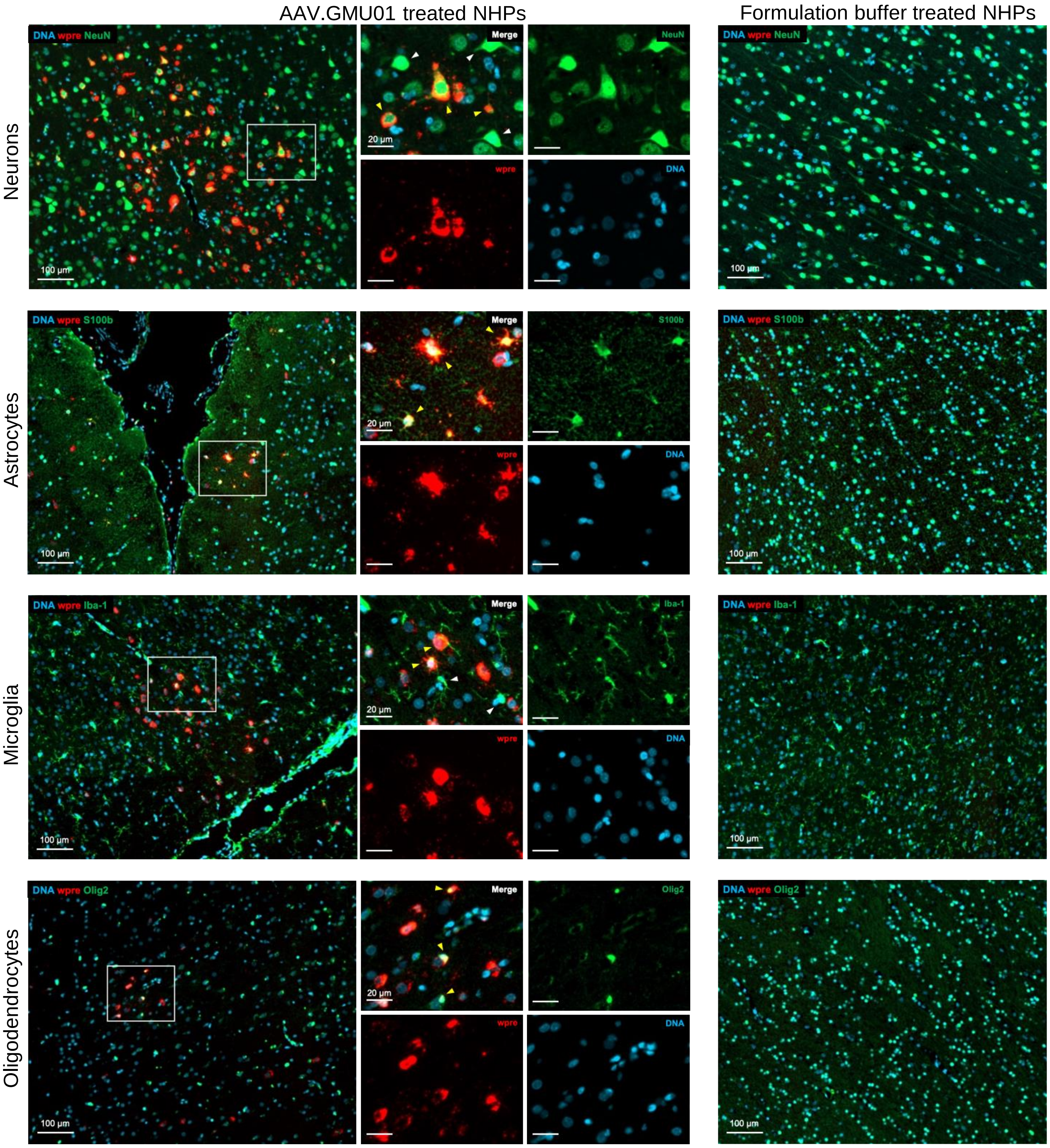
AAV.GMU01 treated NHPs
Formulation buffer treated NHPs
Neurons
Astrocytes
Microglia
Oligodendrocytes
DNA wpre NeuN
DNA wpre S100b
DNA wpre Iba-1
DNA wpre Olig2
Merge
NeuN
S100b
Iba-1
Olig2
wpre
DNA
100 µm
20 µm

**Supplemental Figure 3: AAV.GMU01 shows evidence of transducing neurons, astrocytes, microglia and oligodendrocytes in NHP brain.** Representative images depicting co-detection of WPRE RNA and different cell type markers. Yellow arrowhead: transduced cell; White arrowhead: un-transduced cell. Staining and imaging details in Methods section.

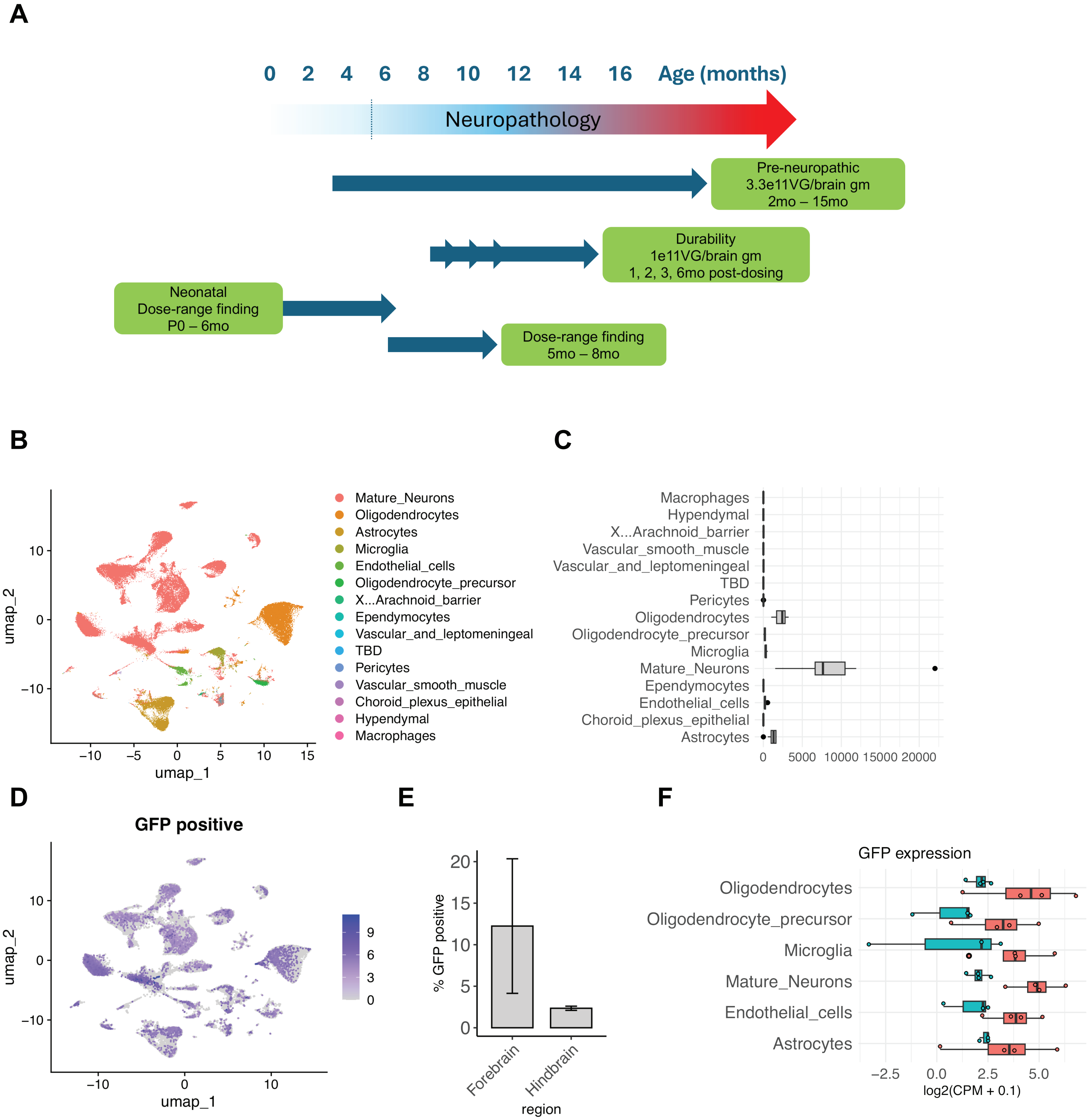
A
0 2 4 6 8 10 12 14 16 Age (months)
Neuropathology
Pre-neuropathic
3.3e11VG/brain gm
2mo – 15mo
Durability
1e11VG/brain gm
1, 2, 3, 6mo post-dosing
Neonatal
Dose-range finding
P0 – 6mo
Dose-range finding
5mo – 8mo
B
umap_1
umap_2
Mature_Neurons
Oligodendrocytes
Astrocytes
Microglia
Endothelial_cells
Oligodendrocyte_precursor
X...Arachnoid_barrier
Ependymocytes
Vascular_and_leptomeningeal
TBD
Pericytes
Vascular_smooth_muscle
Choroid_plexus_epithelial
Hypendymal
Macrophages
C
Macrophages
Hypendymal
X...Arachnoid_barrier
Vascular_smooth_muscle
Vascular_and_leptomeningeal
TBD
Pericytes
Oligodendrocytes
Oligodendrocyte_precursor
Microglia
Mature_Neurons
Ependymocytes
Endothelial_cells
Choroid_plexus_epithelial
Astrocytes
0 5000 10000 15000 20000
D
GFP positive
umap_1
umap_2
E
% GFP positive
Forebrain
Hindbrain
region
F
GFP expression
Oligodendrocytes
Oligodendrocyte_precursor
Microglia
Mature_Neurons
Endothelial_cells
Astrocytes
log2(CPM + 0.1)

**Supplemental Figure 4: Pharmacology studies in *Arsa* KO mice treated with AAV.GMU01-*ARSA*** (**A**) Overview of pharmacology studies in *Arsa* KO mice treated with AAV.GMU01-*ARSA*. Single-cell RNA sequencing in mouse brain treated with AAV.GMU01-GFP (**B**) UMAP of cell states in mouse brains transfected with AAV.GMU01. (**C**) Distribution of cell states in the data. Total cell frequencies (left) and proportions of cell states (right) are presented. (**D**) Feature plot of GFP positive cells on UMAP projection. (**E**) Percentage of GFP positive cells in the forebrain and hindbrain in different samples. Error bars represent standard error of the means between the samples. (**F**) Bar plots of GFP expression per cell states in the forebrain (red bars) and hindbrain (teal bars). Dots represent expression in different samples.

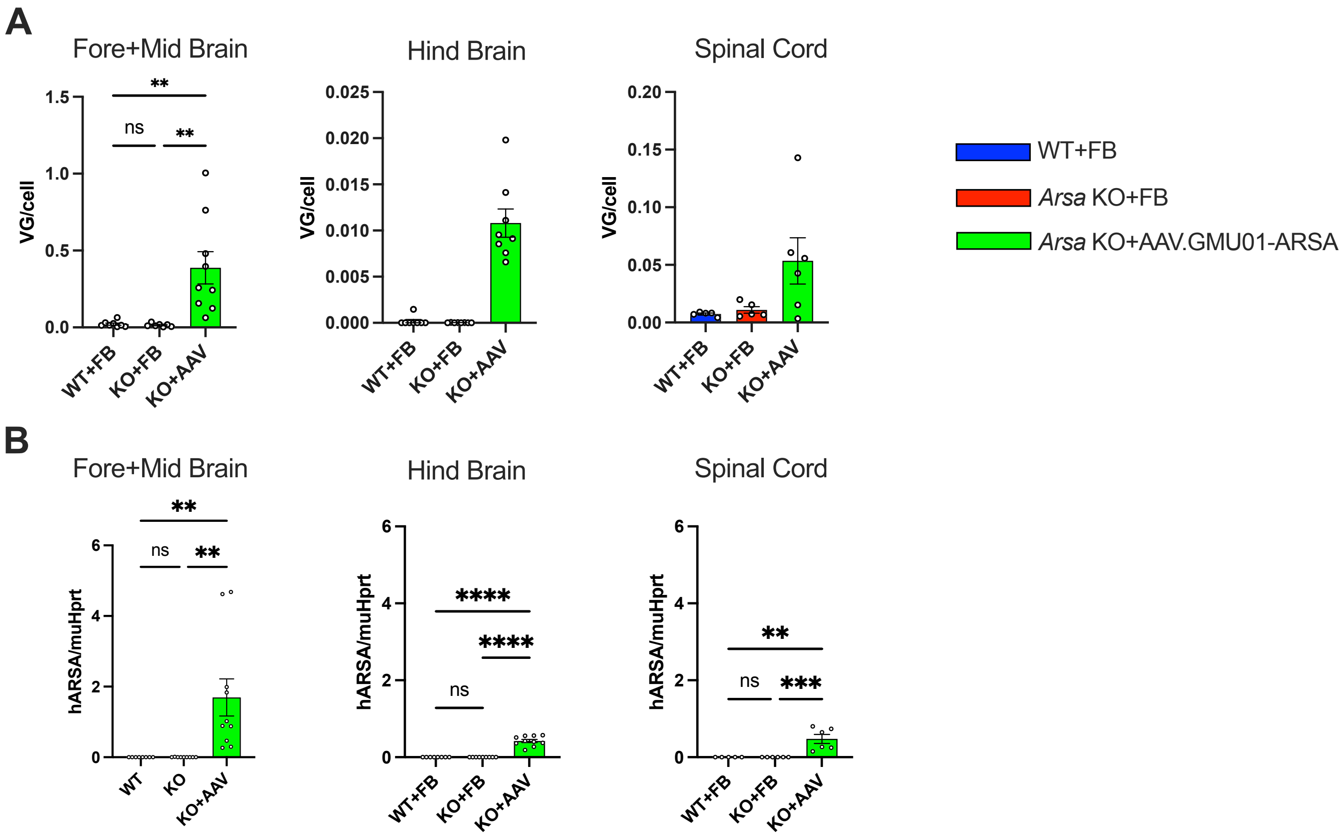

A
Fore+Mid Brain
Hind Brain
Spinal Cord
VG/cell
1.5
1.0
0.5
0.0
0.025
0.020
0.015
0.010
0.005
0.000
0.20
0.15
0.10
0.05
0.00
**
ns
**
WT+FB
KO+FB
KO+AAV
WT+FB
Arsa KO+FB
Arsa KO+AAV.GMU01-ARSA
B
Fore+Mid Brain
Hind Brain
Spinal Cord
hARSA/muHprt
6
4
2
0
**
ns
**
****
****
ns
**
ns
***
WT
KO
KO+AAV
WT+FB
KO+FB
KO+AAV

**Supplemental Figure 5: *Arsa* KO Mice Treated with AAV.GMU01-*ARSA* show CNS-wide ARSA expression.** Pre-neuronopathic *Arsa* KO mice and age-matched control animals were dosed with formulation buffer or AAV.GMU01-*ARSA* at 1.6e11VG/mouse (3.3e11VG/gram brain weight). Thirteen months post-dose, *Arsa* KO mice were euthanized, and brain and spinal cord samples collected. (**A**) Vector exposure by bGH-dPCR, (**B**) human *ARSA* mRNA levels by RT-dPCR. Error bars represent mean with standard error. One-way ANOVA with Tukey's multiple comparison test. *$p$<0.05; **$p$<0.01; ***$p$<0.001; ****$p$<0.0001. FB= Formulation Buffer.

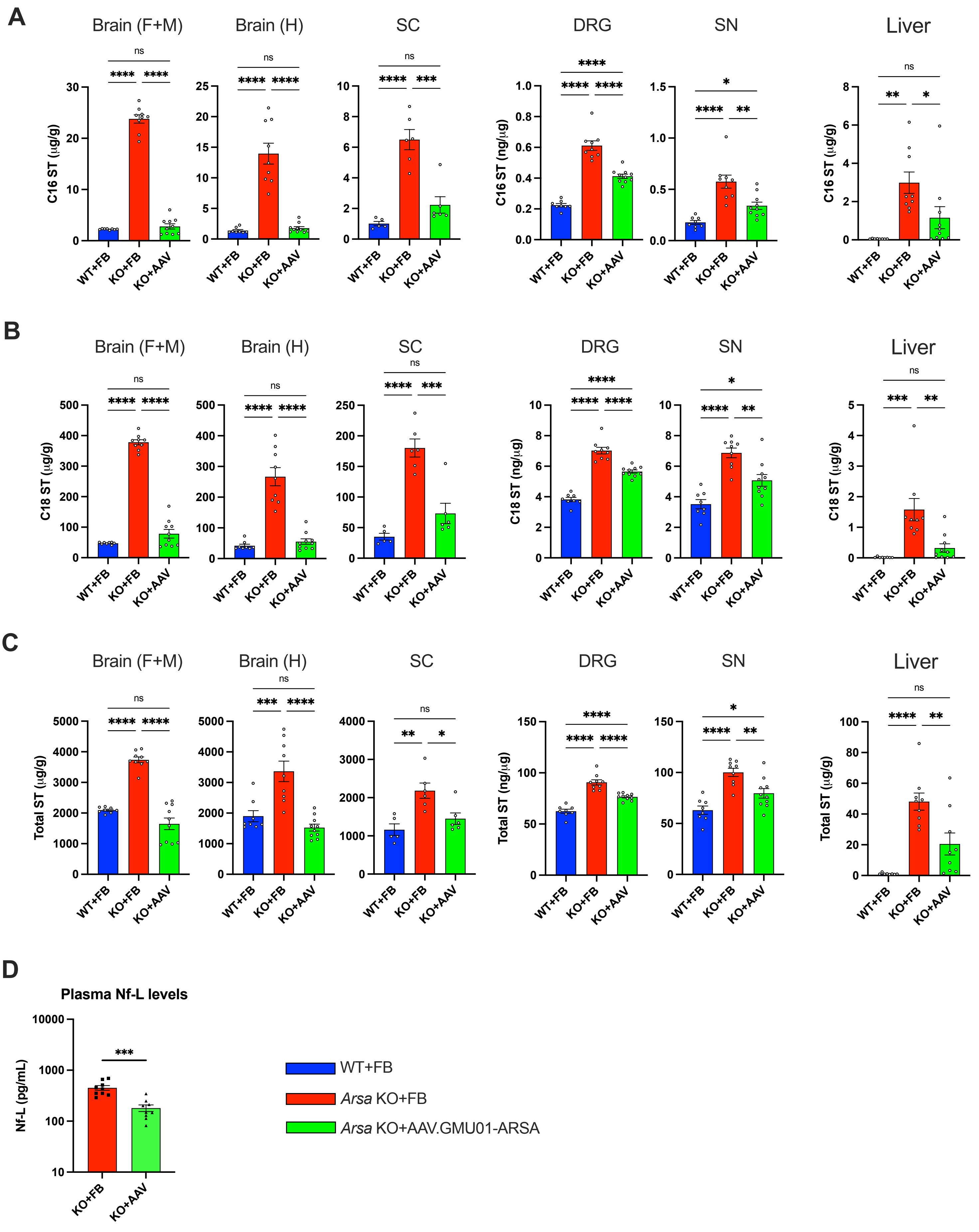
A
Brain (F+M)
Brain (H)
SC
DRG
SN
Liver
C16 ST (μg/g)
C16 ST (ng/μg)
B
C18 ST (μg/g)
C18 ST (ng/μg)
C
Total ST (μg/g)
Total ST (ng/μg)
WT+FB
KO+FB
KO+AAV
ns
D
Plasma Nf-L levels
Nf-L (pg/mL)
WT+FB
Arsa KO+FB
Arsa KO+AAV.GMU01-ARSA

**Supplemental Figure 6: Phenotypic reversal in *Arsa* KO mice treated with AAV.GMU01-*ARSA*.** Pre-neuronopathic *Arsa* KO mice and age-matched control animals were dosed with formulation buffer or with AAV.GMU01-*ARSA* at 1.6e11VG/mouse (3.3e11VG/gram brain weight). Thirteen months post-dose, *Arsa* KO mice were euthanized and brain, spinal cord, DRG, sciatic nerve and liver samples collected. (**A-C**) Sulfatide levels were measured using liquid chromatography–mass spectrometry (LC-MS). Data normalized to tissue weight: converted from ng/mL (50 µL) to µg/g (ng/µg for DRG and SN). **(D)** Plasma samples were collected at necropsy and were assayed by Quanterix Simoa to quantify neurofilament light chain (NfL). Error bars represent mean with standard error. One-way ANOVA with Tukey's multiple comparison test. *$p$<0.05; **$p$<0.01; ***$p$<0.001; ****$p$<0.0001. FB= Formulation Buffer, sulfatide= ST.

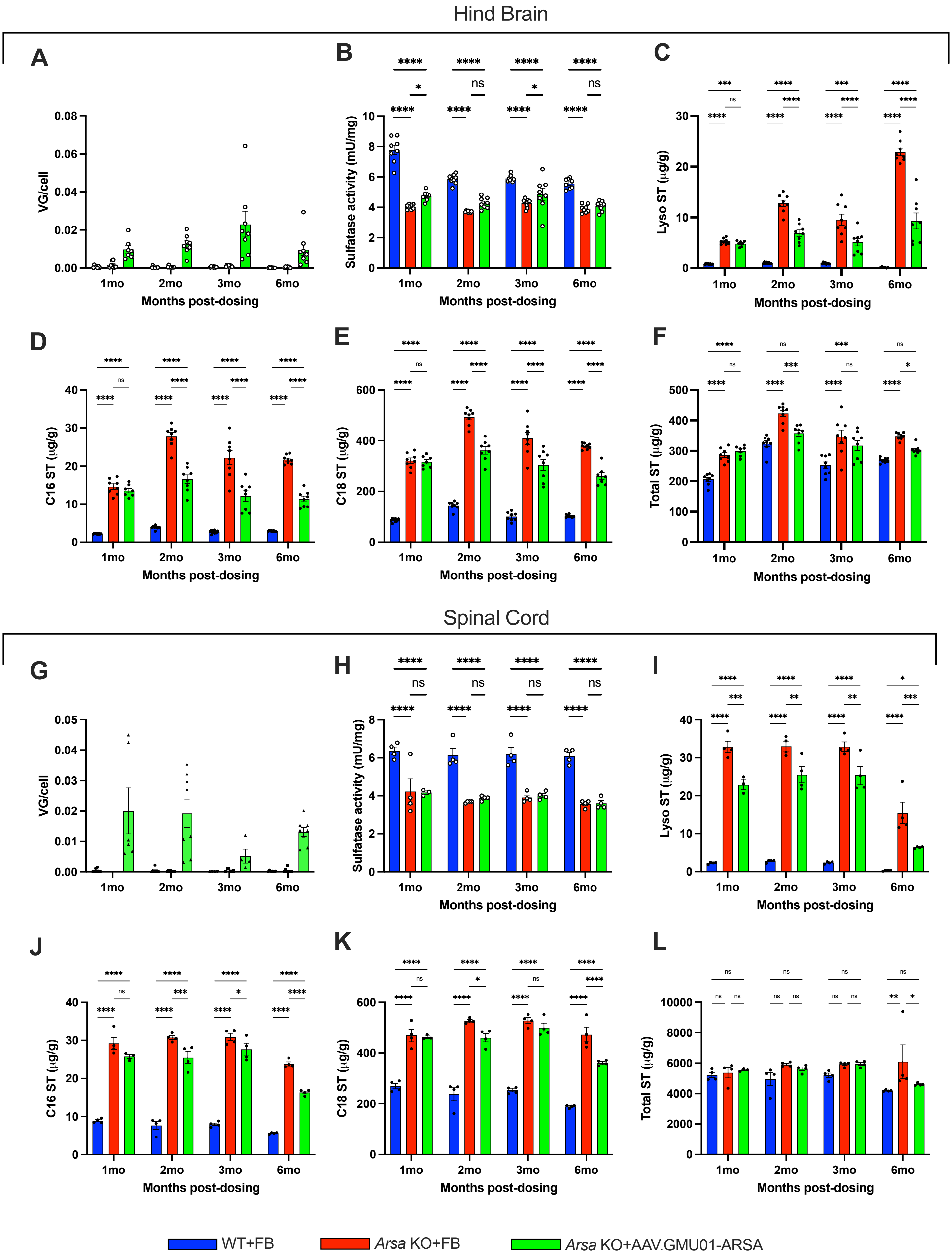

Hind Brain
A
B
C
VG/cell
Sulfatase activity (mU/mg)
Lyso ST (μg/g)
Months post-dosing
1mo
2mo
3mo
6mo
D
E
F
C16 ST (μg/g)
C18 ST (μg/g)
Total ST (μg/g)
Spinal Cord
G
H
I
VG/cell
Sulfatase activity (mU/mg)
Lyso ST (μg/g)
J
K
L
C16 ST (μg/g)
C18 ST (μg/g)
Total ST (μg/g)
WT+FB
Arsa KO+FB
Arsa KO+AAV.GMU01-ARSA

**Supplemental Figure 7: ARSA expression and function is persistent over time.** Early-neuronopathic *Arsa* KO mice (6mo at dosing) and age-matched control animals were dosed with formulation buffer or AAV.GMU01-*ARSA* at 5e10VG/mouse (1.0e11VG/gram brain weight). One, two, three and six months post-dose, mice were euthanized, and samples collected. (**A-F**) Hind brain samples, (**G-L**) Spinal Cord. (**A, G**) Vector exposure by bGH-dPCR normalized to the *Rab1a* gene (intronic region). (**B, H**) ARSA-mediated sulfatase activity was measured using the Sulfatase Activity Assay Kit, data normalized to total protein measured by BCA assay. (**C, I**) Lyso-sulfatide, (**D, J**) C16-sulfatide isoform, (**E, K**) C18-sulfatide isoform and (**F, L**) Total sulfatide levels were measured using LC-MS. Data normalized to tissue weight: converted from ng/mL (50 µL) to µg/g. Error bars represent mean with standard error. Two-way ANOVA with Tukey's multiple comparison test. *$p$<0.05; **$p$<0.01; ***$p$<0.001; ****$p$<0.0001. FB= Formulation Buffer, sulfatide= ST.

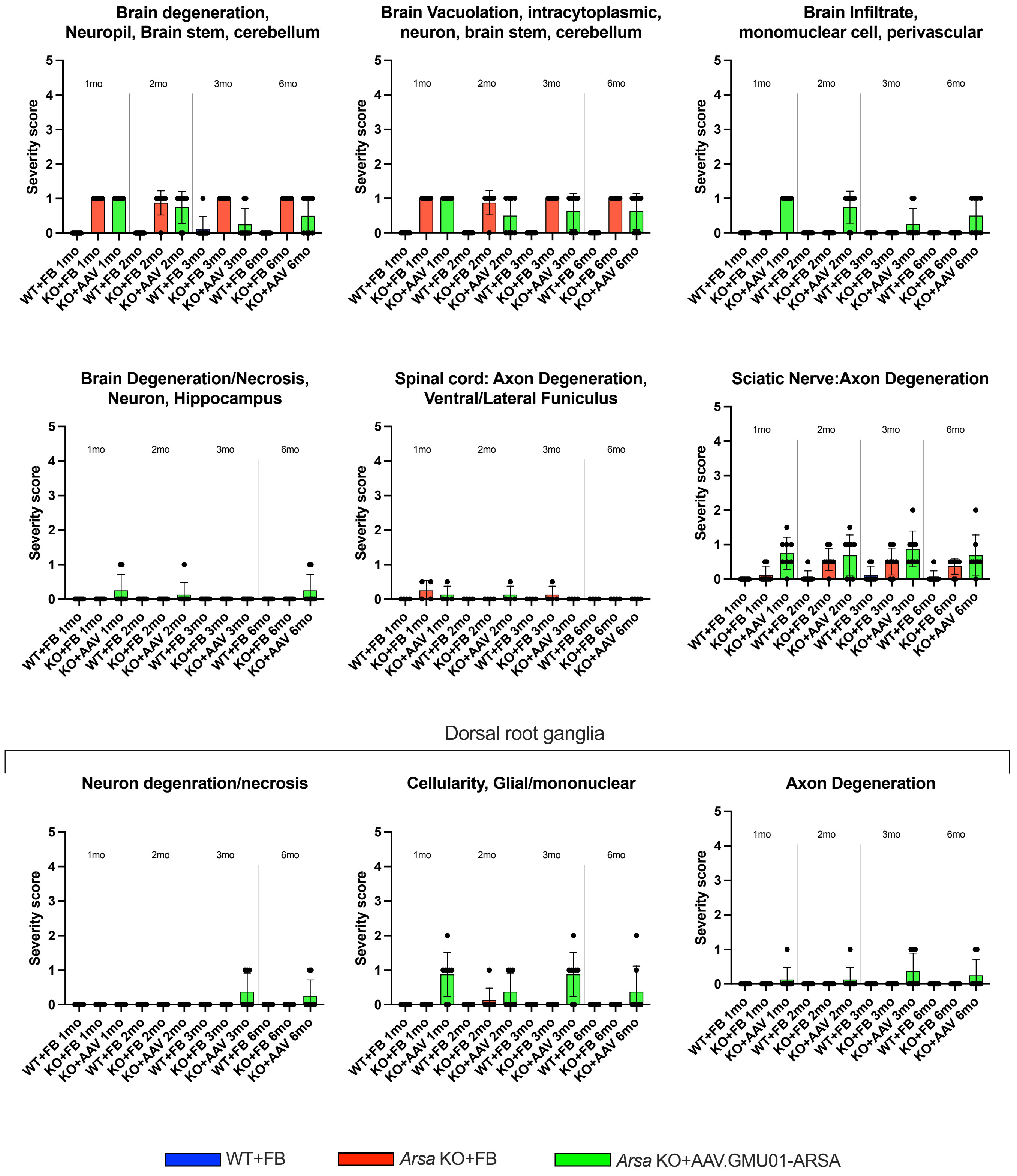

Brain degeneration, Neuropil, Brain stem, cerebellum
Brain Vacuolation, intracytoplasmic, neuron, brain stem, cerebellum
Brain Infiltrate, monomuclear cell, perivascular
Brain Degeneration/Necrosis, Neuron, Hippocampus
Spinal cord: Axon Degeneration, Ventral/Lateral Funiculus
Sciatic Nerve:Axon Degeneration
Dorsal root ganglia
Neuron degenration/necrosis
Cellularity, Glial/mononuclear
Axon Degeneration
Severity score
1mo
2mo
3mo
6mo
WT+FB 1mo
KO+FB 1mo
KO+AAV 1mo
WT+FB 2mo
KO+FB 2mo
KO+AAV 2mo
WT+FB 3mo
KO+FB 3mo
KO+AAV 3mo
WT+FB 6mo
KO+FB 6mo
KO+AAV 6mo
WT+FB
Arsa KO+FB
Arsa KO+AAV.GMU01-ARSA

**Supplemental Figure 8: AAV.GMU01-*ARSA* treated *Arsa* KO mice show up to mild findings.** Histopathological findings in brain, spinal cord, sciatic nerve and DRG in *Arsa* KO treated at 6 months (early-neuronopathic); each data point represents the maximum severity of findings scored on 1-2 sections per animal. Severity scores refer to findings graded as 0= no findings, 1=minimal, 2=mild, 3=moderate, 4=marked, 5=severe. Error bars represent mean with standard error. FB= Formulation Buffer.

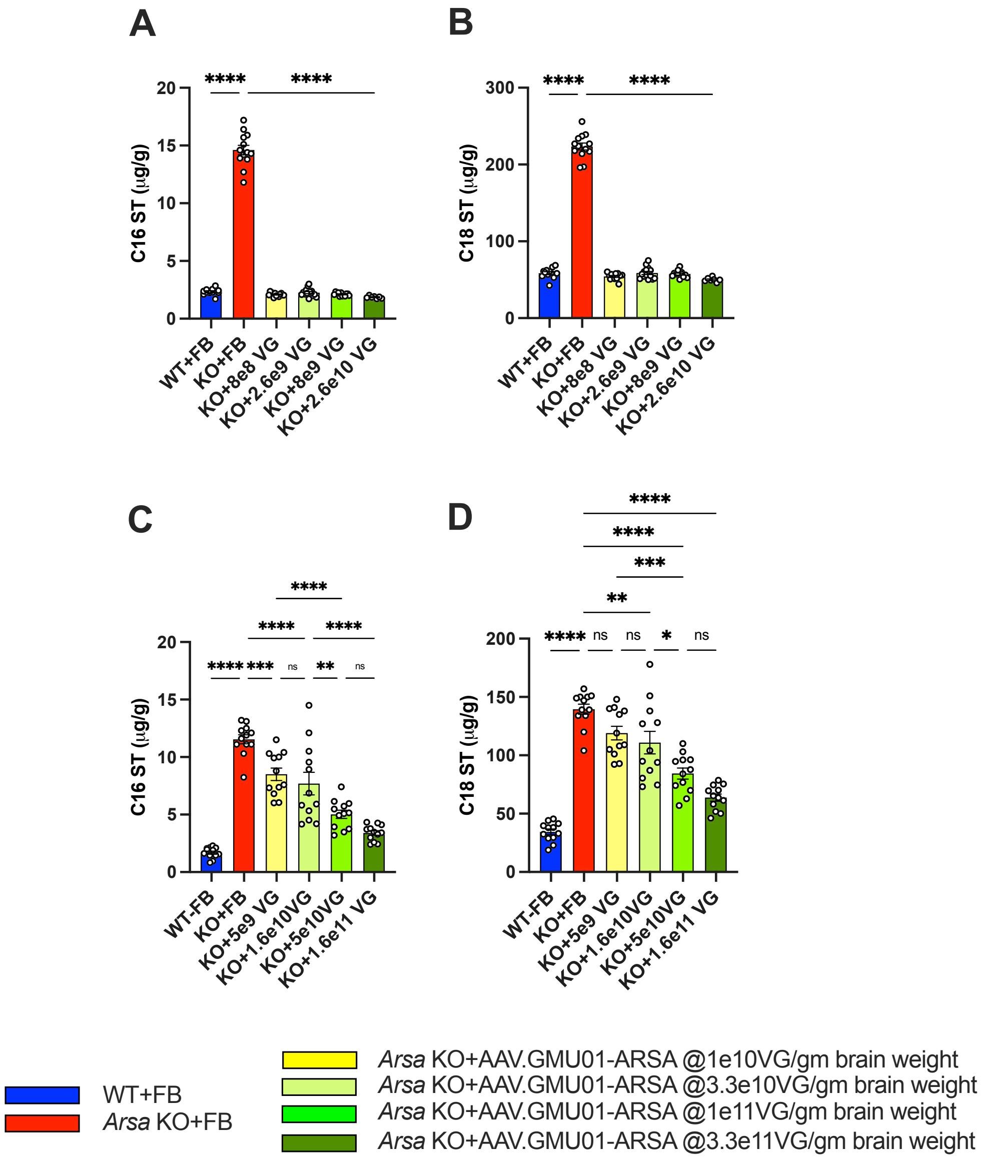
A
C16 ST (µg/g)
WT+FB
KO+FB
KO+8e8 VG
KO+2.6e9 VG
KO+8e9 VG
KO+2.6e10 VG
B
C18 ST (µg/g)
C
WT-FB
KO+5e9 VG
KO+1.6e10VG
KO+5e10VG
KO+1.6e11 VG
D
WT+FB
Arsa KO+FB
Arsa KO+AAV.GMU01-ARSA @1e10VG/gm brain weight
Arsa KO+AAV.GMU01-ARSA @3.3e10VG/gm brain weight
Arsa KO+AAV.GMU01-ARSA @1e11VG/gm brain weight
Arsa KO+AAV.GMU01-ARSA @3.3e11VG/gm brain weight

**Supplemental Figure 9. AAV.GMU01-*ARSA* treated results in dose-dependent sulfatide clearance in *Arsa* KO mice.** (**A-B**) Neonatal *Arsa* KO mice (P0 at dosing) and age-matched control animals were dosed with formulation buffer or AAV.GMU01-*ARSA* at the given doses. Six months post-dose, mice were euthanized, and samples collected. (**C-D**) Early-neuronopathic *Arsa* KO mice (6mo at dosing) and age-matched control animals were dosed with AAV.GMU01-*ARSA* at noted doses. Three months post-dose, mice were euthanized, and samples collected. (**A, C**) C16 sulfatide isoform and (**B, D**) C18 sulfatide isoform levels were measured using LC-MS. Data normalized to tissue weight: converted from ng/mL (50 µL) to µg/g. Data normalized to tissue weight: converted from ng/mL (50 µL) to µg/g. Error bars represent mean with standard error. Two-way ANOVA with Tukey's multiple comparison test. *$p$<0.05; **$p$<0.01; ***$p$<0.001; ****$p$<0.0001.  FB= Formulation Buffer, sulfatide= ST; gbw= grams per brain weight.

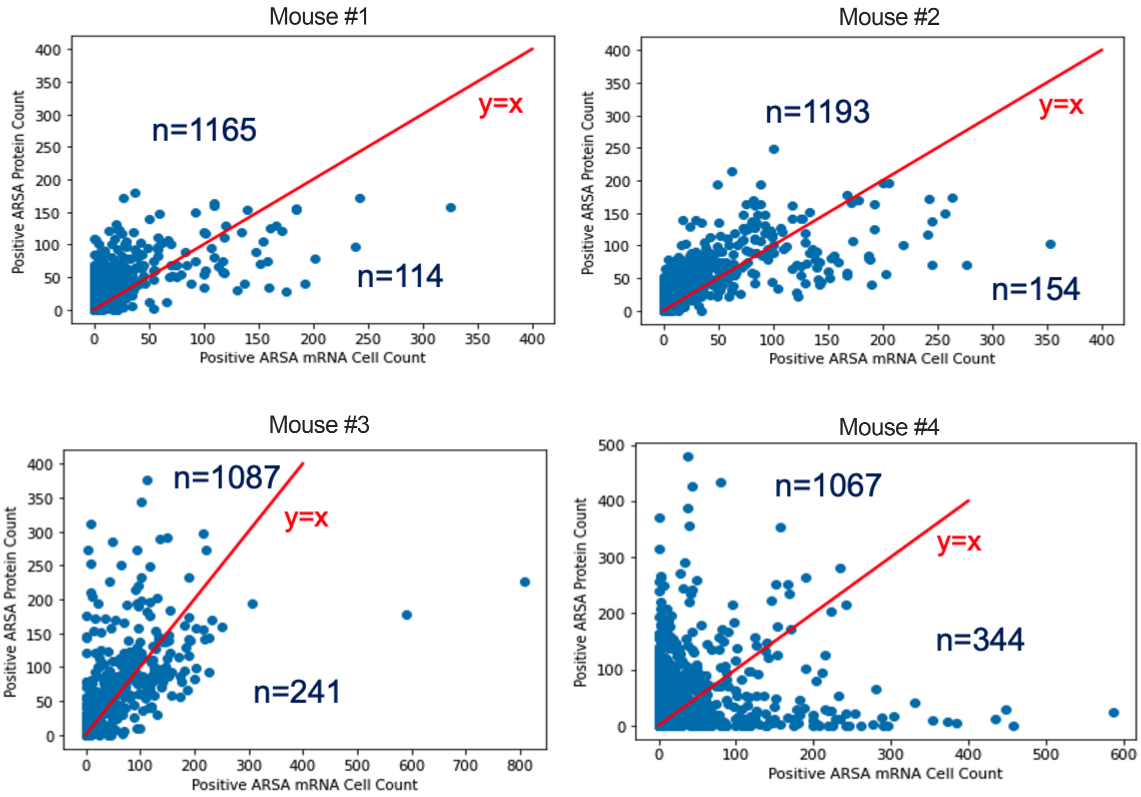
Mouse #1
n=1165
y=x
n=114
Positive ARSA Protein Count
Positive ARSA mRNA Cell Count
Mouse #2
n=1193
y=x
n=154
Positive ARSA Protein Count
Positive ARSA mRNA Cell Count
Mouse #3
n=1087
y=x
n=241
Positive ARSA Protein Count
Positive ARSA mRNA Cell Count
Mouse #4
n=1067
y=x
n=344
Positive ARSA Protein Count
Positive ARSA mRNA Cell Count

**Supplemental Figure 10: AAV.rh10-CBA-ARSA-WPRE treated *Arsa* KO mice show evidence of ARSA protein cross-correction.** Late-stage (13 month) *Arsa* KO mice were dosed with AAV.rh10-CBA-ARSA-WPRE. Three months post-dosing, ARSA-mRNA *in situ* hybridization (ISH) and ARSA-protein immunohistochemistry (IHC) were performed on matched sagittal brain hemi-sections. The sections were imaged and analyzed for signal overlay. Adjacent sagittal sections from mouse brain were treated for WPRE ISH or ARSA IHC with DAPI staining for nuclei. The sagittal sections were then analyzed as individual 1024x1024 pixel tiles. In each tile, ISH positive cells and IHC positive cells were determined by thresholding parameters empirically determined across the entire sagittal images. ISH+ cell count vs IHC+ cell count from each tile is plotted as a scatter plot with y=x line shown in red. Tiles above the y=x line indicates cross-corrected cells.

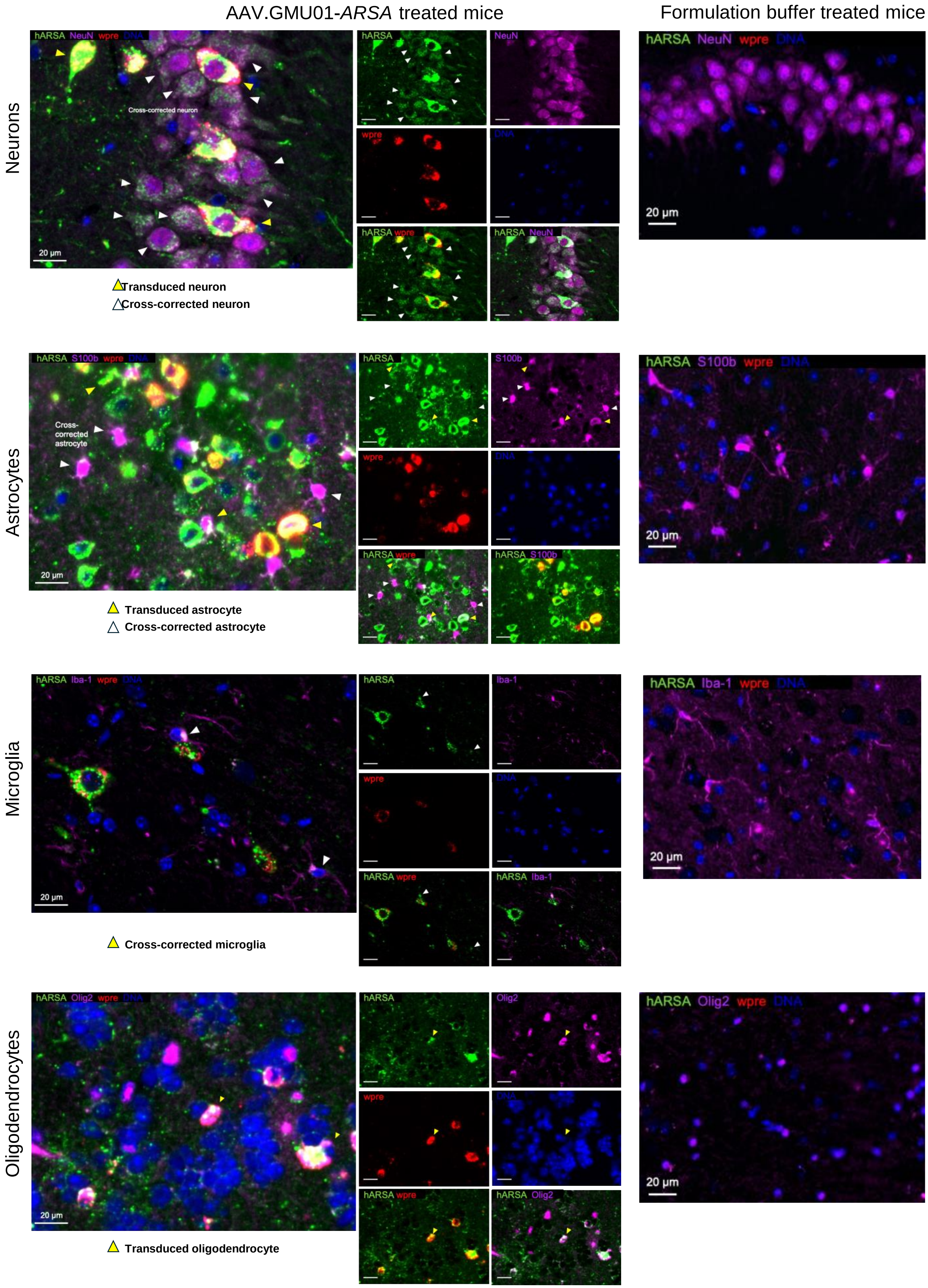
AAV.GMU01-ARSA treated mice
Formulation buffer treated mice
Neurons
hARSA NeuN wpre DNA
Cross-corrected neuron
20 µm
hARSA
NeuN
wpre
DNA
hARSA wpre
hARSA NeuN
Transduced neuron
Cross-corrected neuron
Astrocytes
hARSA S100b wpre DNA
Cross-corrected astrocyte
S100b
hARSA S100b
Transduced astrocyte
Cross-corrected astrocyte
Microglia
hARSA Iba-1 wpre DNA
Iba-1
hARSA Iba-1
Cross-corrected microglia
Oligodendrocytes
hARSA Olig2 wpre DNA
Olig2
hARSA Olig2
Transduced oligodendrocyte

**Supplemental Figure 11: Evidence of ARSA protein cross-correction in *Arsa* KO mice upon treatment with AAV.GMU01-*ARSA*.** Representative images depicting co-detection of WPRE RNA, ARSA protein and different cell type markers. Yellow arrowhead: transduced cell (positive for wpre and cell-type marker); White arrowhead: cross-corrected cell (positive for ARSA and cell-type marker but negative for wpre). Staining and imaging details in Methods section.

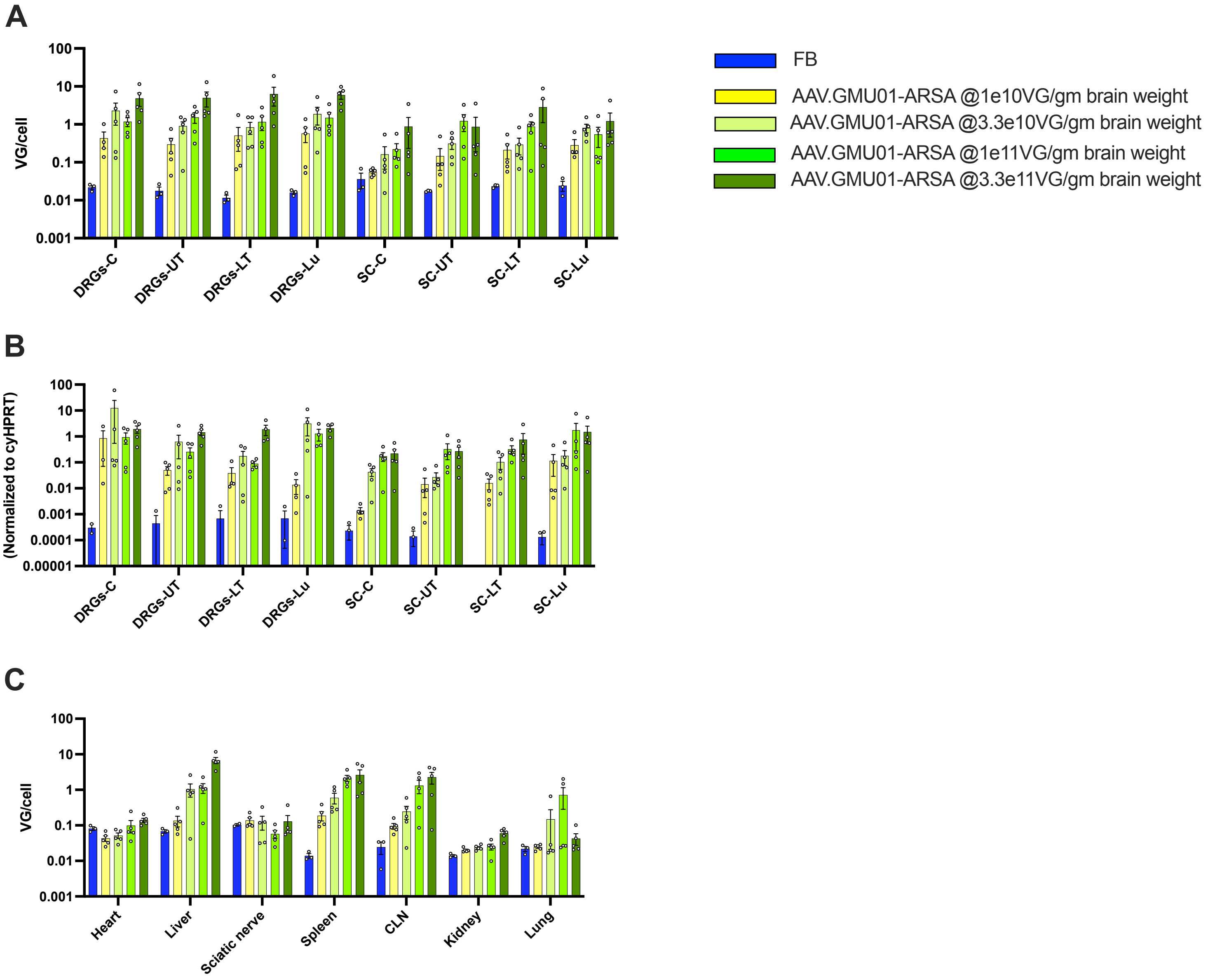
A
B
C
FB
AAV.GMU01-ARSA @1e10VG/gm brain weight
AAV.GMU01-ARSA @3.3e10VG/gm brain weight
AAV.GMU01-ARSA @1e11VG/gm brain weight
AAV.GMU01-ARSA @3.3e11VG/gm brain weight
VG/cell
100
10
1
0.1
0.01
0.001
DRGs-C
DRGs-UT
DRGs-LT
DRGs-Lu
SC-C
SC-UT
SC-LT
SC-Lu
hARSA mRNA
(Normalized to cyHPRT)
0.0001
0.00001
Heart
Liver
Sciatic nerve
Spleen
CLN
Kidney
Lung

**Supplemental Figure 12: Widespread dose-dependent vector biodistribution and ARSA expression in spinal cord, DRGs and peripheral tissues.** Eight spinal cord segments with adjacent DRGs, 2 each from cervical, upper thoracic, lower thoracic and lumbar were flash frozen and DNA/RNA isolated. (**A, C**) Digital PCR (dPCR) was performed to quantify AAV.GMU01 vector concentration and normalized to the *TUBB1* gene intron. (**B**) RT-dPCR was performed to quantify *ARSA* mRNA expression, normalized to endogenous *HPRT* gene. Each data point represents VG/cell exposure or normalized *ARSA* expression for that sample, averaged across all NHPs in that group. Error bars represent mean with standard error. FB= Formulation Buffer. C=cervical, UT=upper thoracic; LT=lower thoracic; Lu=lumbar; CLN=cervical lymph node

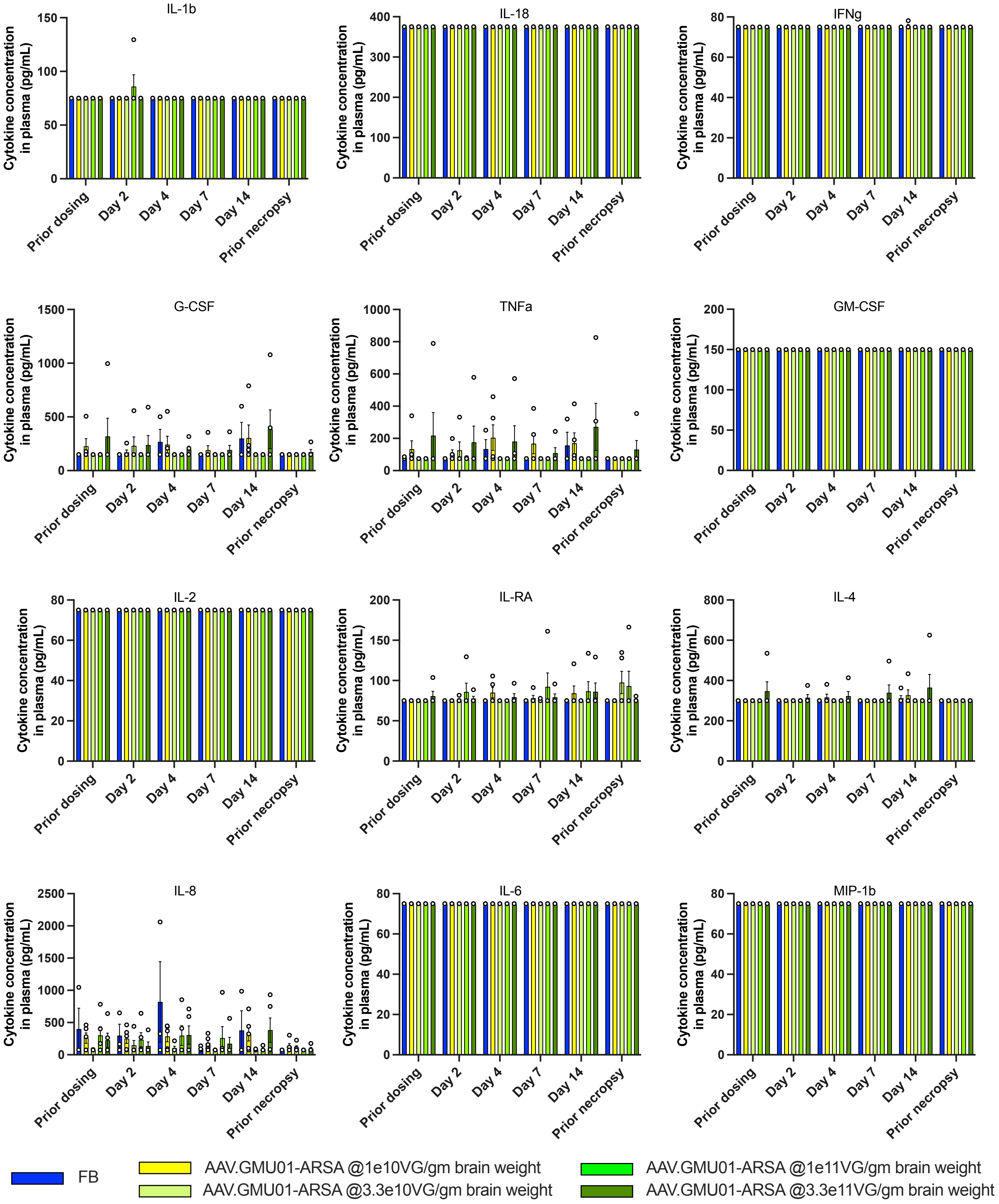
IL-1b
IL-18
IFNg
G-CSF
TNFa
GM-CSF
IL-2
IL-RA
IL-4
IL-8
IL-6
MIP-1b
Cytokine concentration in plasma (pg/mL)
Prior dosing
Day 2
Day 4
Day 7
Day 14
Prior necropsy
FB
AAV.GMU01-ARSA @1e10VG/gm brain weight
AAV.GMU01-ARSA @3.3e10VG/gm brain weight
AAV.GMU01-ARSA @1e11VG/gm brain weight
AAV.GMU01-ARSA @3.3e11VG/gm brain weight

**Supplemental Figure 13: ICM infusion was well tolerated and did not trigger innate immune responses.** Plasma was isolated pre-dose and at Days 2, -4, 7, 14 post-dose and at necropsy. The Luminex assay was used to determine the concentration of IL-1b, IL-1RA, IL-6, IL-10, IL12/23 (p40), IL-15, IL-18, IFN-g, TNF-a, G-CSF, MCP-1, MIP-1b, GM-CSF, IL-2, IL-4, IL-5, IL-8, IL-13 and IL-17A. Each data point represented cytokine concentration in that sample, averaged across all NHPs in that group. Error bars represent mean with standard error.

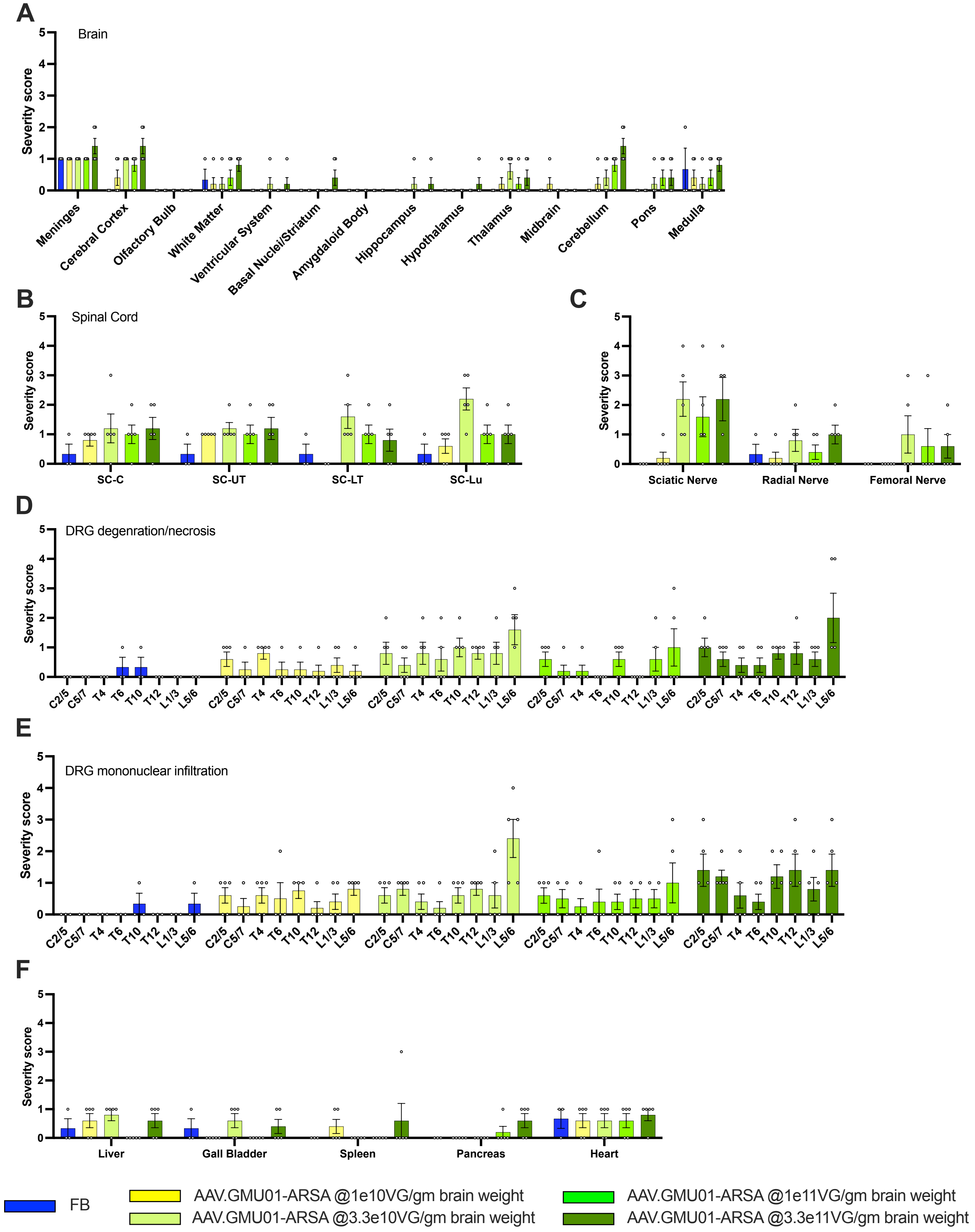

A
Brain
Severity score
Meninges
Cerebral Cortex
Olfactory Bulb
White Matter
Ventricular System
Basal Nuclei/Striatum
Amygdaloid Body
Hippocampus
Hypothalamus
Thalamus
Midbrain
Cerebellum
Pons
Medulla
B
Spinal Cord
SC-C
SC-UT
SC-LT
SC-Lu
C
Sciatic Nerve
Radial Nerve
Femoral Nerve
D
DRG degenration/necrosis
C2/5
C5/7
T4
T6
T10
T12
L1/3
L5/6
E
DRG mononuclear infiltration
F
Liver
Gall Bladder
Spleen
Pancreas
Heart
FB
AAV.GMU01-ARSA @1e10VG/gm brain weight
AAV.GMU01-ARSA @3.3e10VG/gm brain weight
AAV.GMU01-ARSA @1e11VG/gm brain weight
AAV.GMU01-ARSA @3.3e11VG/gm brain weight

**Supplemental Figure 14: ICM infusion was well tolerated in NHPs.** Histopathological findings in brain, spinal cord, and DRG; each data point represents the maximum severity of findings scored on 1-2 sections per animal. Refer to Supplementary Table 6 for study design. Severity scores refer to findings graded as 0= no findings, 1=minimal, 2=mild, 3=moderate, 4=marked, 5=severe. Error bars represent mean with standard error. C=cervical, T=thoracic; L=lumbar (numbers represent segments reviewed by pathologist)

### SUPPLEMENTAL TABLES

#### Supplemental Table 1: Study design evaluating AAV.GMU01 and AAV.rh10 vector biodistribution and transgene expression in CNS

| Group | # of Animals | Test Article | Dose (vg) | Dose (vg/gm brain weight) | RoA | Dosing paradigm | Necropsy |
|---|---|---|---|---|---|---|---|
| 1 | 3 (M) | Formulation Buffer | N/A | N/A | Intrathecal Catheter (C1-C2) | 2.5ml>6hr >2.5ml | 29 days |
| 2 | 3 (M) | AAV.rh10-CBA-eGFP | 2.75E+13 | 2.65E+11 | | | 16 days |
| 3 | 3 (M) | AAV.GMU01-eGFP | 2.75E+13 | 2.65E+11 | | | 18 days |

Internal

**Supplemental Table 2: Study design comparing AAV.GMU01 biodistribution and transgene activity by ICV and ICM administration routes**

| Group | Animal #’ | ROA | Description | Animal position | Dose | Volume | Rate | Necropsy |
|---|---|---|---|---|---|---|---|---|
| 1 | 3 (M) | Bilateral ICV | Clearpoint, Sequential | Sternal recumbent followed by Trendelenburg | 2.0e13 VG/NHP | 1ml per hemisphere | 0.125ml/min | 28±1 days |
| 2 | 3 (M) | Direct ICM | CM puncture with needle | Trendelenburg | | 2ml | | |

**Supplemental Table 3: Long-term Pharmacology, Efficacy and Durability Study in *Arsa* KO mice (2mo-15mo)**

| Group | Number/sex | Genotype | Test article | Dose | Dosing regimen | Time Points |
|---|---|---|---|---|---|---|
| **1** | 12 (6M, 6F) | WT | Formulation buffer | N/A | Bi-lateral ICV, 4 uL each side | Dosing age: 2 mo Necropsy age: 15 mo (13 mo in-life) |
| **2** | 12 (6M, 6F) | $ARSA^{-/-}$ | Formulation buffer | N/A | | |
| **3** | 12 (6M, 6F) | $ARSA^{-/-}$ | AAV.GMU01-*ARSA* | 1.6e11VG (3.3e11VG/gram brain weight) | | |

**Supplemental Table 4: Longitudinal Pharmacology, Efficacy and Durability Study in *Arsa* KO Mice**

| Group | Number and Sex | Genotype | Test Article | Dose | Dosing Regimen | Time Points |
|---|---|---|---|---|---|---|
| **1** | 32 (16M, 16F) | WT | Formulation buffer | N/A | Bi-lateral ICV, 4ul each side | Dosing age: 6 mo<br>Necropsy age: 7 mo (1 mo in-life), 8 mo (2 mo in-life), 9 mo (3 mo in-life), 12 mo (6 mo in-life) |
| **2** | 32 (16M, 16F) | *Arsa*$^{-/-}$ | Formulation buffer | N/A | | |
| **3** | 32 (16M, 16F) | *Arsa*$^{-/-}$ | AAV.GMU01-*ARSA* | 5e10VG (1e11VG/gram brain weight) | | |

Internal

### Supplemental Table 5: Dose-dependent pharmacology studies in *Arsa* KO Mice

<table>
<tr><th colspan="7">Pharmacology and dose optimization study in P0 neonatal mice</th></tr>
<tr><td>Group</td><td>Number and Sex</td><td>Genotype</td><td>Test Article</td><td>Dose</td><td>Dosing Regimen</td><td>Time Points</td></tr>
<tr><td>1</td><td>12 (6M, 6F)</td><td>WT</td><td>Formulation buffer</td><td>N/A</td><td rowspan="6">Bi-lateral ICV, 3ul each side</td><td rowspan="6">Dosing age: P0<br>Necropsy: 6 mo<br>(6mo in-life)</td></tr>
<tr><td>2</td><td>12 (6M, 6F)</td><td rowspan="5">Arsa$^{-/-}$</td><td>Formulation buffer</td><td>N/A</td></tr>
<tr><td>3</td><td>12 (6M, 6F)</td><td rowspan="4">AAV.GMU01-ARSA</td><td>2.64e10VG (3.3e11VG/gram brain weight)</td></tr>
<tr><td>4</td><td>12 (6M, 6F)</td><td>8.0e9VG (1.0e11VG/gram brain weight)</td></tr>
<tr><td>5</td><td>12 (6M, 6F)</td><td>2.64e9VG (3.3e10VG/gram brain weight)</td></tr>
<tr><td>6</td><td>12 (6M, 6F)</td><td>8.0e8VG (1.0e10VG/gram brain weight)</td></tr>
<tr><td></td><td></td><td></td><td></td><td></td><td></td><td></td></tr>
<tr><th colspan="7">Pharmacology and dose optimization study in early-neuronopathic mice</th></tr>
<tr><th>Group</th><th>Number and Sex</th><th>Genotype</th><th>Test Article</th><th>Dose</th><th>Dosing Regimen</th><th>Time Points, Sample Collection, Analysis</th></tr>
<tr><td>1</td><td>12 (6M, 6F)</td><td>WT</td><td>Formulation buffer</td><td>N/A</td><td rowspan="6">Bi-lateral ICV, 5ul each side</td><td rowspan="6">Dosing age: 6 mo<br>Necropsy: 9 mo<br>(3mo in-life)</td></tr>
<tr><td>2</td><td>12 (6M, 6F)</td><td rowspan="5">Arsa$^{-/-}$</td><td>Formulation buffer</td><td>N/A</td></tr>
<tr><td>3</td><td>12 (6M, 6F)</td><td rowspan="4">AAV.GMU01-ARSA</td><td>1.6e11VG (3.3e11VG/gram brain weight)</td></tr>
<tr><td>4</td><td>12 (6M, 6F)</td><td>- 5.0e10VG (1.0e11VG/gram brain weight)</td></tr>
<tr><td>5</td><td>12 (6M, 6F)</td><td>- 1.6e10VG (3.3e10VG/gram brain weight)</td></tr>
<tr><td>6</td><td>12 (6M, 6F)</td><td>- 5.0e9VG (1.0e10VG/gram brain weight)</td></tr>
</table>

### Supplemental Table 6: NHP dose-ranging and tolerability study

<table>
<tr><th>Group</th><th>Test Article</th><th>Dose/gm Brain Weight</th><th>Dose/Animal</th><th>Dosing Regimen</th><th>Animal #’s</th><th>Time Points, Sample Collection</th></tr>
<tr><td>1</td><td rowspan="4">AAV.GMU01-ARSA</td><td>1e10 VG</td><td>7.5e11 VG</td><td rowspan="5">RoA: ICM with animal placed in Trendelenburg position<br><br>Doing parameters: 2.5ml @0.125mL/min with 250ul flush</td><td>5 (M/F)</td><td rowspan="5">In-life: 35 days<br><br>CSF (pre-study, and at necropsy)<br><br>Plasma: pre-study, days post-dose: 2, 4, 7, 14 and at necropsy<br><br>Neurological/behavioral assessment (pre-dose, 7 days and 5 weeks post-dose)</td></tr>
<tr><td>2</td><td>3.3e10 VG</td><td>2.5e12 VG</td><td>5 (M/F)</td></tr>
<tr><td>3</td><td>1e11 VG</td><td>7.5e12 VG</td><td>5 (M/F)</td></tr>
<tr><td>4</td><td>3.3e11 VG</td><td>2.5e13 VG</td><td>5 (M/F)</td></tr>
<tr><td>5</td><td>Formulation Buffer</td><td>n/a</td><td>n/a</td><td>3 (M/F)</td></tr>
</table>

**Supplemental Table 7. Neurological and Behavioral Battery performed pre-dosing, on day 7 post-dose and at necropsy (Week 5 Post-dose).**

| General observations | Attitudinal Posture Reactions | Spinal Segmental Reflex | Cranial Nerve Function |
|---|---|---|---|
| Mental status | Tonic neck | Perineal Reflex | Pathologic Nystagmus |
| Head posture | Hopping | Withdrawal reflex | Facial sensory Exam |
| Motor activity | Visual placing | Cross extensor | Olfactory Nerve Exam |
| Coordination | Tactile placing | Patellar reflex | Jaw tone |
| Stance | Postural reaction | Babinski reflex | Gag reflex |
| Gait | | Chaddock reflex | Facial symmetry |
| Circling | | | Cochlear Nerve Exam |
| | | | Tongue Exam |
| | | | Menace Response |
| | | | Pupilliary light reflex (PLR) |
| | | | PLR Consensual |
| | | | Pupil Size |
| | | | Pupil symmetry |
| | | | Ocular position |
| | | | Ocular motility |
| | | | Oculovestibular reflex |
| | | | Palpebral reflex |
